# ROS responsive Aux/IAA multimerization modulates auxin responses

**DOI:** 10.1101/2024.02.12.579961

**Authors:** Dipan Roy, Poonam Mehra, Vaishnavi Mukkawar, Lisa Clark, Kevin Bellande, Joop EM Vermeer, Raquel Martin Arevallilo, Teva Vernoux, Kawinnat Sue ob, Andrew Jones, Ulrike Bechtold, Phil Mullineaux, Kathryn Lilley, Adrian Brown, Malcolm Bennett, Ari Sadanandom

## Abstract

Reactive oxygen species (ROS) function as key signals in plants to enable adaptation to environmental stresses. Plant roots respond to transient water stress by temporarily ceasing branching using the acclimative response xerobranching^1^. In this study, we report that a rapid ROS burst regulates Xerobranching by inducing multimerization of auxin repressor protein IAA3/SHY2. Mutations in specific cysteine residues in IAA3/SHY2 disrupt redox-mediated multimerization and interaction with co-repressor TPL, but not with auxin response partner ARF7 and auxin receptor TIR1. ROS-mediated oligomerization of IAA3/SHY2 is required for efficient ARF mediated target gene repression during Xerobranching and lateral root emergence. We demonstrate that AUX/IAA proteins vary in their redox mediated multimerization, revealing a new auxin response regulatory mechanism that directly connects ROS sensing to auxin signalling. Our study reveals how ROS, auxin and water stress intersect to shape acclimative responses in plant roots and maintain their phenotypic plasticity.

---

Plants are exposed to a myriad of environmental stresses that trigger acclimative responses. Plant cells use ROS as an “alarm” signal for environmental stresses that functions to coordinate adaptive responses^2^. To test whether the acclimative response Xerobranching (XB) is dependent on ROS, roots of Arabidopsis wildtype and ROS mutants were exposed to a Xerobranching bioassay^3^. Roots of *Arabidopsis* RBOH (Respiratory Burst Oxidase Homolog) mutants continued to branch when growing across an air-gap, in contrast to wildtype controls **(Figure 1a, b)**. Hence, ROS functions as a key regulatory signal during a Xerobranching response.

**Figure 1.**
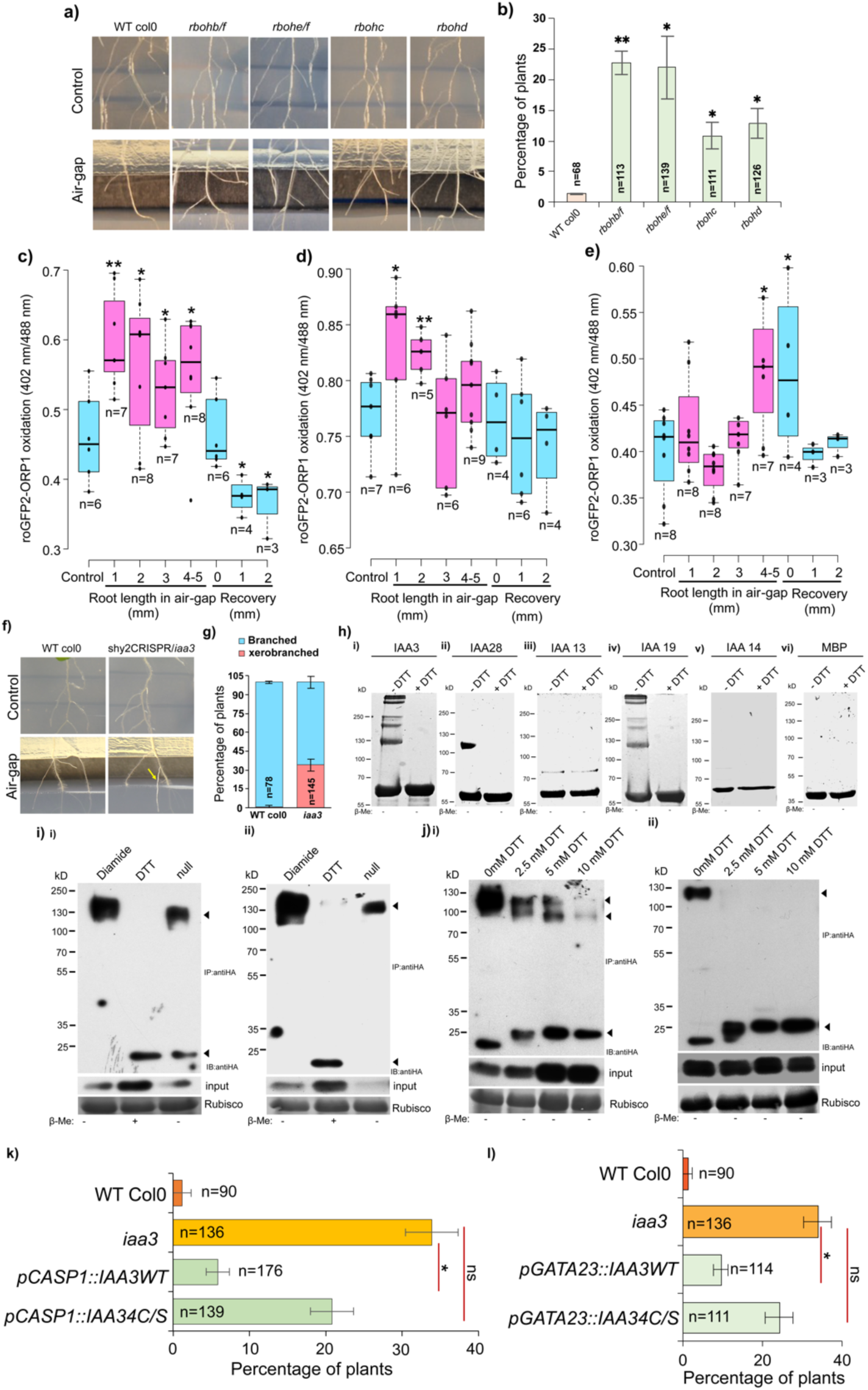
AUX/IAA repressor IAA3 /SHY2 regulates Xerobranching by sensing changes in cellular redox status via selected cysteine residues. (a,b) Agar based air-gap assay system showing xerobranching response in Arabidopsis thaliana (air gap ∼5mm). Representative images (a) and a bar graph (b) of agar-based air gap assay showing xerobranching defect in *rboh* mutants compared to WT col0. (c-e) Xerobranching stimulus triggers ROS production in Arabidopsis root tips exposed to air-gap. ROS sensor, roGFP2-orp1 shows Hydrogen peroxide levels in (c) nucleus, (d) cytoplasm, and (e)apoplast. Numbers in horizontal axis indicate length of root tips grown into air-gap and subsequent recovery conditions. (f)CRISPR/Cas9 knockout line of Arabidopsis IAA3/SHY2 (referred interchangeably as shy2 CRISPR or iaa3) disrupt xerobranching unlike WT Col0. (g) Frequency plot showing percentage of branched and xerobranched WT Col 0 and *iaa3* plants in air-gaps. (h) Redox modulated cysteine disulphide linkage dependent multimerization of candidate Aux/IAAs (i)IAA3 (ii)IAA28 (iii) IAA13 (iv) IAA19 (v) IAA14 (vi) MBP (Maltose Binding Protein) tag control (i) Redox dependent multimerization pattern of constitutively expressed WT-IAA3/SHY2 (left panel) and 4C/S-IAA3/SHY2 (right panel) in *Arabidopsis thaliana*. (j) DTT concentration-based titration assays indicating the decrease in the propensity of 4C/S-IAA3/SHY2 (right panel) in forming redox dependent multimers compared to WT IAA3/SHY2 (left panel). (k,l) pCASP1 and pGATA23 driven expression of 4C/S-IAA3/SHY2 cannot complement the xerobranching defect of iaa3 plants in comparison to WT-IAA3/SHY2. Bars represent percentage of plants showing xerobranching defect. Statistical significance was calculated using Student’s *t*-test*, *p≤0.05,* ** *p≤0.01,* n.s. ‘not significant’ differences.

To identify *<u>where</u>* and *<u>when</u>* the ROS burst occurs following a Xerobranching stimulus, we employed versions of the H_2_O_2_ sensor roGFP2-orp1^4^ which are either targeted to cytoplasmic, apoplastic or nuclear subcellular compartments^5^. Reduced roGFP2-orp1 proteins display a characteristic single peak at ∼490 nm excitation, whilst the oxidised protein exhibits two peaks at 400 nm and 490 nm, thereby allowing quantitative assessment of redox states through the ratio of fluorescence intensity between 400 ±5 nm and 490 ±5 nm. For example, seedlings expressing the nuclear localized nls-roGFP2-orp1 H_2_O_2_ sensor display a significant increase in 402/488 ratio when growing root tips traverses across an air gap in the Xerobranching bioassay. This indicates nls-roGFP2-orp1 to be more oxidised due to a ROS burst in the nuclei of root cells exposed to transient water stress **(Figure 1c-1e)**. However, once root tips reconnect with agar (recovery phase), nuclear ROS levels fall rapidly. Temporal analysis of nls-roGFP2-orp1 oxidation state revealed an increase in nuclear ROS levels within 4-5 hours after a Xerobranching stimulus **(Figure 1c)**. In contrast to other subcellular compartments, in root cell nuclei, ROS burst was maintained for the duration of the Xerobranching response.

_W_hat are the nuclear targets that ROS modify to trigger a Xerobranching response? Earlier work reported the Xerobranching response pathway acts by modifying the re-distribution of the root branching hormone auxin^3^. This led us to consider whether the Xerobranching-induced nuclear ROS burst modifies components of the auxin response machinery. To pinpoint ARF (Auxin Response Factor) and/or AUX/IAA auxin response components required for Xerobranching, we characterised *loss of function* mutations in root expressed members of these families using our bioassay. Our phenotypic analysis revealed that the *AUX/IAA* gene *IAA3/SHY2* was functionally important for the Xerobranching response. We generated an *Arabidopsis* CRISPR/cas9 knockout line disrupting *IAA3/SHY2*^6^, with deletions encompassing part of the *SHY2* promoter, 5’ UTR and exon 1 and exon 2 (alleles C1 and C2). Our preliminary phenotypic analysis indicated that both allele C1 and C2 had a lateral root phenotype (**Supplementary Figure S2)** and all our subsequent analysis were taken forward with the larger deletion harbouring allele C1 (*shy2* CRISPR line henceforth). Our xerobranching assays indicated that CRISPR/cas9 mediated disruption of *IAA3/SHY2* caused ∼ 40% of the *shy2* CRISPR seedlings to branch in air gaps **(Figure 1f-1g)**. Hence, our results reveal the importance of the repressive function of IAA3/SHY2 in orchestrating a Xerobranching response.

How may the nuclear ROS burst regulate Xerobranching via AUX/IAAs like IAA3/SHY2? It has been hypothesised that oligomerization of Aux/IAAs may be required for efficient ARF mediated repression of targets^7,8^. However, how this Aux/IAA oligomerization might be triggered, and the underpinning mechanism remains elusive. We hypothesise that cysteine disulphide linkage mediated multimerization of the Aux/IAAs in a redox dependent manner could provide a mechanism for efficient ARF mediated repression in plant nuclei. To test this hypothesis, we first assessed the distribution and conservation of cysteine residues amongst Aux/IAAs **(Supplementary figure S1a)**. We observed widespread divergence in the frequency and distribution of cysteine residues with only cysteine 171 residue conserved amongst Aux/IAAs. Next, we explored whether redox conditions influence Aux/IAA multimerization. We selected candidate Aux/IAAs that fulfilled the following criteria: (i) representative of the different clades of Aux/IAAs (ii) exhibit a wide spectrum of cysteine residues (iii) have well defined regulatory functions (iv) with roles during lateral root development. The candidate Aux/IAAs were recombinantly expressed and purified as N-terminal MBP (maltose binding protein) tagged proteins in non-reducing conditions and profiled by SDS polyacrylamide gel electrophoresis (PAGE). The results revealed that in non-reducing conditions, selected Aux/IAAs like IAA3/SHY2 and IAA19/MSG2 formed higher order multimers **(Figure 1h)**, and other Aux/IAAs, like IAA28, IAA14 and IAA13 were largely monomeric **(Figure 1h)**. Moreover, subsequent assays with redox modifying agents also indicated that selected Aux/IAAs like IAA3/SHY2 and IAA19/MSG2 are more sensitive to redox changes and forms highly robust higher order multimers (**Supplementary figure S3**), indicating that the ability of Aux/IAAs like IAA3/SHY2 to form higher order redox-dependent multimers provides a mechanism to vary auxin-dependent transcriptional outputs in response to environmental cues that are modulated through dynamic changes in cellular redox status.

Our multimerization assays reveal that IAA3/SHY2 is a highly ROS responsive Aux/IAA **(Figure 1h and Supplementary figure S3)**. How does IAA3/SHY2 redox dependent multimerization affect its repressive function? In order to assess this, we first identified the cysteine residues that facilitate IAA3/SHY2 homo-multimer formation. Since multimerization is postulated to occur through the Aux/IAA PB1 domains^9–11^, we reasoned that cysteine residues in this domain may confer ROS responsive multimerization of IAA3/SHY2. Cysteine residues 112, 169, 171 and 187 of IAA3/SHY2 are part of the PB1 domain (**Supplementary figure S1a-1b**), while cysteine residues 23, 31 and 32 are between domains DI and DII^12^, indicating that sulfhydryl bridges mediated by cysteine residues localized in the PB1 domain could stabilise IAA3/SHY2 multimers. To test the effect of manipulating these cysteine residues on multimerization of IAA3/SHY2, we generated cysteine to serine substitution mutations. The Cys169Ser mutation of IAA3/SHY2 resulted in the most significant loss of higher order multimers, while Cys32Ser and Cys112Ser mutations resulted in a drastic reduction of trimeric and tetrameric forms of IAA3/SHY2. Cys187Ser mutation in IAA3/SHY2 resulted in slight reduction of almost all the higher order multimeric forms of the protein with a significant reciprocal increase in the monomeric form under non-reducing conditions (**Supplementary figure S4a).** Quadruple Cys32Ser, Cys112Ser, Cys169Ser and Cys187Ser mutations in IAA3/SHY2 (henceforth as IAA3/SHY2-4C/S or 4C/S-IAA3/SHY2) blocked formation of multimers in non-reducing **(Supplementary figure S4b)** and oxidizing **(Supplementary figure S 4c)** conditions in vitro. While 7.5mM DTT reduced IAA3-WT to monomers, but it only took treatment with the oxidising agent diamide at 6mM concentration to reform the multimers. Whilst IAA3-4C/S was monomeric even at 15mM diamide failing to form any multimers **(Supplementary figure S4c (ii))**. To determine which cysteine residues are more responsive to redox changes and are more likely to be involved in inter- and intra-molecular disulphide bond formation in IAA3-WT, we modified the native protein with N-ethylmaleimide (NEM), followed by reduction and alkylation with DTT and iodoacetamide respectively. We then carried out trypsin digestion followed by LC-MS/MS to identify which cysteine residues were modified with NEM and which with iodoacetamide. Our rationale was that natively reduced cysteines free to be labelled with NEM were less ROS responsive and thus less likely to be engaged in disulphide bonds, whereas significantly more ROS responsive cysteines potentially forming disulphide bridges would be labelled with iodoacetamide. Our Mass spectrometry-based analysis revealed that Cys 32, Cys112, Cys169, Cys171 and Cys187 form disulphide bridges as they were labelled with iodoacetamide and not NEM **(Table 1)**, indicating that these cysteine residues were more ROS responsive and were potentially engaged in disulphide bridge formation. Cys23 and Cys112 were observed in both the NEM and iodoacetamide modified forms. This could be indicative of regions of the IAA3/SHY2 structure that are only partially involved in disulphide formation and represent more flexible regions of multimers. This data clearly supports the observations of the significant loss of multimerisation of the Cys169Sser, Cys32Sser, and Cys112Ser, Cys187ser and IAA3-4C/S mutated forms of IAA3-WT **(Supplementary Figure 4)**. Hence, IAA3-WT must multimerize through disulphide linkages between preferred cysteine residues in a redox dependent manner *in vitro* with Cys 169 and Cys 32 being the most preferred, followed by Cys 23 and Cys112. Both wildtype and quadruple cysteine mutant forms were expressed *in planta* as N-terminal HA-tagged WT IAA3 and IAA3-4C/S variants in the *shy2* CRISPR background. IAA3-WT retained a significant fraction of higher order multimers even at 5mM DTT, and these multimers could still be observed even at 10mM DTT concentration **(Figure 1i (i) and (ii)**. However, IAA3-4C/S almost completely lost the ability to form higher order multimers even at the lowest concentration of DTT (2.5mM) tested (**Figure 1j (i) and (ii**)). These results were in agreement with what we observed in *Nicotiana benthamiana* system, where IAA3-WT retains higher order multimers at a high concentration of 10mM DTT **(Supplementary figure S5)**. Our results reveal IAA3/SHY2 multimerizes through disulphide linkages involving specific cysteine residues beyond those previously predicted in the PB1 domain.

**Table. 1.**
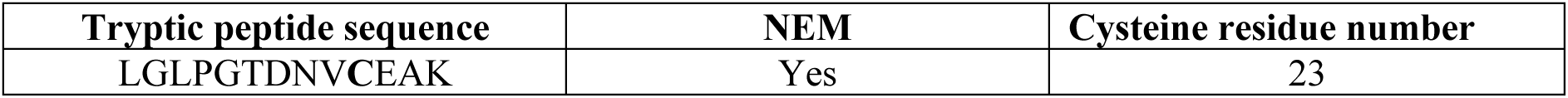
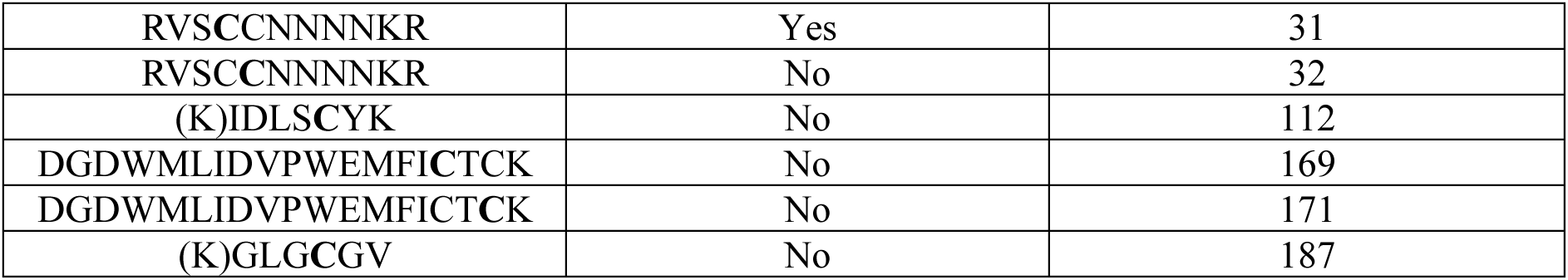
Identification of ROS responsive cysteines of WT-IAA3/SHY2 by Mass spectrometry. Cysteine residues of the native protein that were not labelled by NEM were labelled with carbamidomethyl group post reduction with DT

Next, we tested whether disruption of key cysteine residues for multimerization causes the repressive function of IAA3/SHY2 to be compromised. To test this hypothesis at the phenotypic level, we assessed whether IAA-WT and IAA3-4C/S could rescue the *shy2* CRISPR xerobranching (XB) defect **(Figure 1k-l**). IAA3-WT and IAA3-4C/S was expressed using the CASP1 promoter^13^ and GATA23 promoter^14^ (henceforth as pCASP1::IAA3-WT, pCASP1::IAA3-4C/S, pGATA23::IAA3-WT and pGATA23::IAA3-4C/S) in the *shy2* CRISPR background. We specifically chose the CASP1 promoter because of the distinct lateral root phenotype that has been previously observed when the stabilized form of SHY2, *shy2-2*, is expressed under the control of the CASP1 promoter^13^. GATA23 is a critical component of the auxin signalling module involving IAA28 and ARF5,6,7,8,19 and is required for founder cell identity establishment during lateral root development^14^. We observed that in contrast to pCASP1::IAA3-WT and pGATA23::IAA3-WT, significantly higher frequency of both pCASP1::IAA3-4C/S and pGATA23::IAA3-4C/S seedlings formed lateral roots in the air gap (**Figure 1k-1l**). Hence, Xerobranching assays indicate disrupting redox dependent IAA3/SHY2 multimerization attenuates its repressive function.

Phenotyping transgenic lines for auxin-dependent responses revealed IAA3-4C/S mutants have altered lateral root emergence phenotype opposite to those previously observed for stabilized form of shy2^13^. In our phenotyping assays we observed that pCASP1::IAA3-4C/S exhibit a 2-fold increase in emerged LRs compared to pCASP1::IAA3-WT **(Figure 2B and 2C)**. Hence, endodermal expression of multimer defective IAA3/SHY2-4C/S cannot restore repression by IAA3/SHY2 during LR development. Redox dependent multimerization assays performed with transgenic lines revealed WT-IAA3/SHY2 form more robust redox modulated higher order multimers than 4C/S-IAA3/SHY2 **(Supplementary figure S7)**. Moreover, the repressive effects of IAA3 WT on LR development could not be phenocopied by IAA3-4C/S even when it was being expressed under the control of the 35S promoter^15,16^ (**Supplementary figure S8**) indicating that ROS responsive IAA3 multimer formation is more important for lR repression than IAA3 protein levels per se.

**Figure 2.**
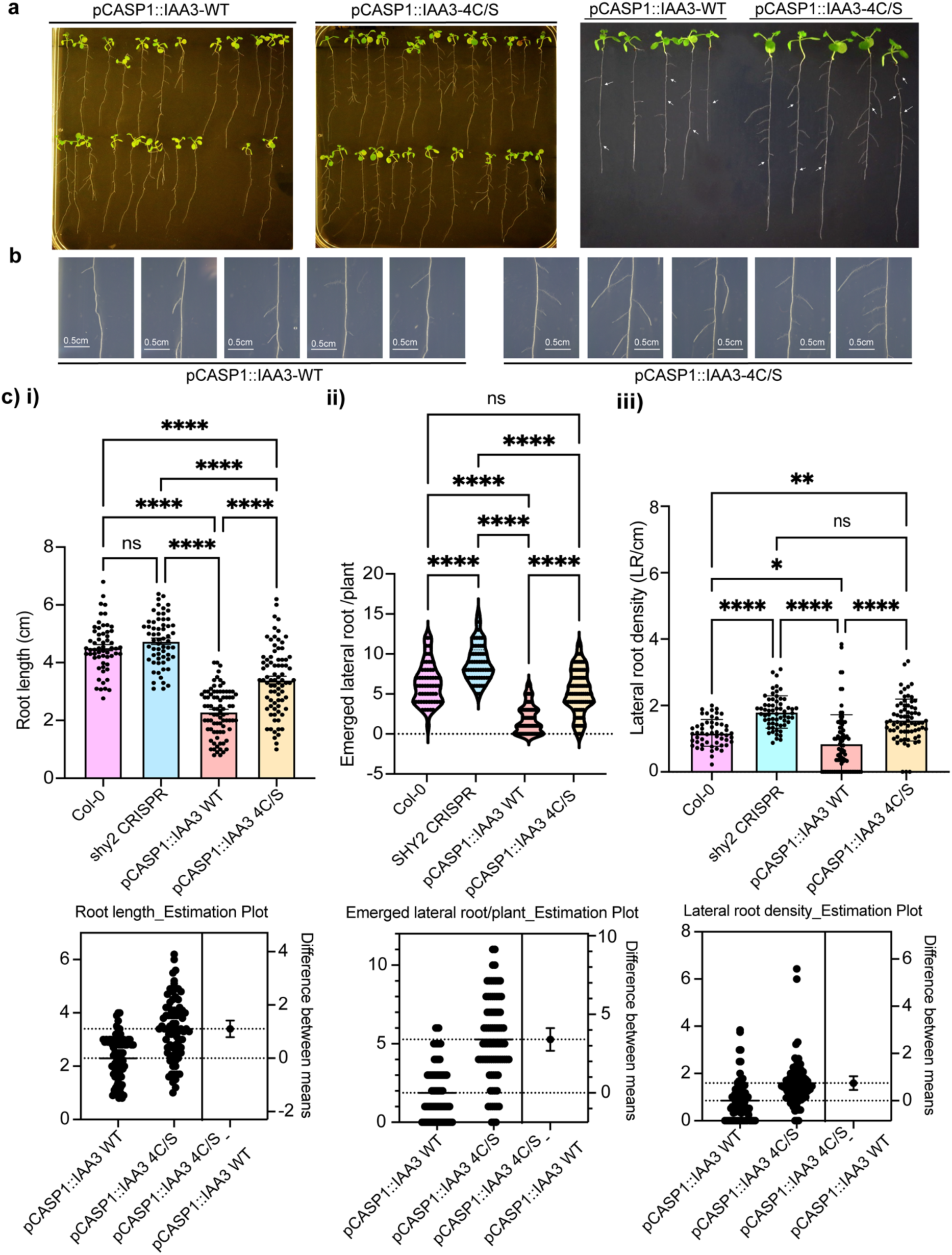
IAA3/SHY2 requires key cysteines involved in multimerization to repress lateral root emergence. Lateral Root (LR) phenotype of pCASP1::IAA3-WT and pCASP1::IAA3-4C/S expressing transgenic seedlings (a,b) Representative images highlighting the difference in the number of emerged lateral roots per plant between pCASP1::IAA3-WTand pCASP1::IAA3-4C/S lines **(c)** Comparative analysis of the differences in (i) root length,(ii) number of emerged lateral roots per plant,(iii) lateral root density between WT col0, shy2 CRISPR, pCASP1::IAA3-WT and pCASP1::IAA3-4C/S lines. ns, not significant, *P ≤ 0.05;***P≤ 0.001 ****P ≤ 0.0001.

GATA23 promoter is expressed specifically in xylem pole pericycle cells before the first asymmetric division and is correlated with oscillating auxin signaling maxima in the basal meristem and is reported to be activated prior to lateral root initiation^14^. Therefore, we hypothesized that expressing IAA3-WT under the control of the GATA23 promoter in the *shy* 2 CRISPR background provides for an engineered closed-loop gene regulatory system, where co-activation of the GATA23 promoter in a manner that is correlated with the oscillating auxin signaling maxima at the basal meristem will lead to steady state levels of the IAA3-WT protein, which in turn will provide a negative feedback signal within the modular auxin signalling network^17^ and repress the process of LR development from its earliest stage. We observed in pGATA23::IAA3WT seedlings a significant reduction in numbers of emerged LRs per plant and lateral root density in comparison to WT Col-0 and *shy2* CRISPR seedlings, whereas there was no significant difference in pGATA23::IAA3-4C/S seedlings **(Figure 3)**. This aberrant temporal retardation in the process of LR development in pGATA23::IAA3-WT seedlings likely reflects the importance of repressive effect of IAA3/SHY2 very early in LR development. In contrast, the absence of a LR phenotype in pGATA23:: IAA3-4C/S seedlings reveals that regulation of redox mediated IAA multimers is critical during the onset of founder cell identity establishment in this developmental program. Indeed, the inability of IAA3-4C/S to repress LR development irrespective of being expressed under CASP1, 35S or GATA23 promoters sheds new light on the importance of how formation of redox dependent multimers provides an alternative route to repress target gene expression for Aux/IAAs like IAA3 in a tissue specific and temporal manner.

**Figure 3.**
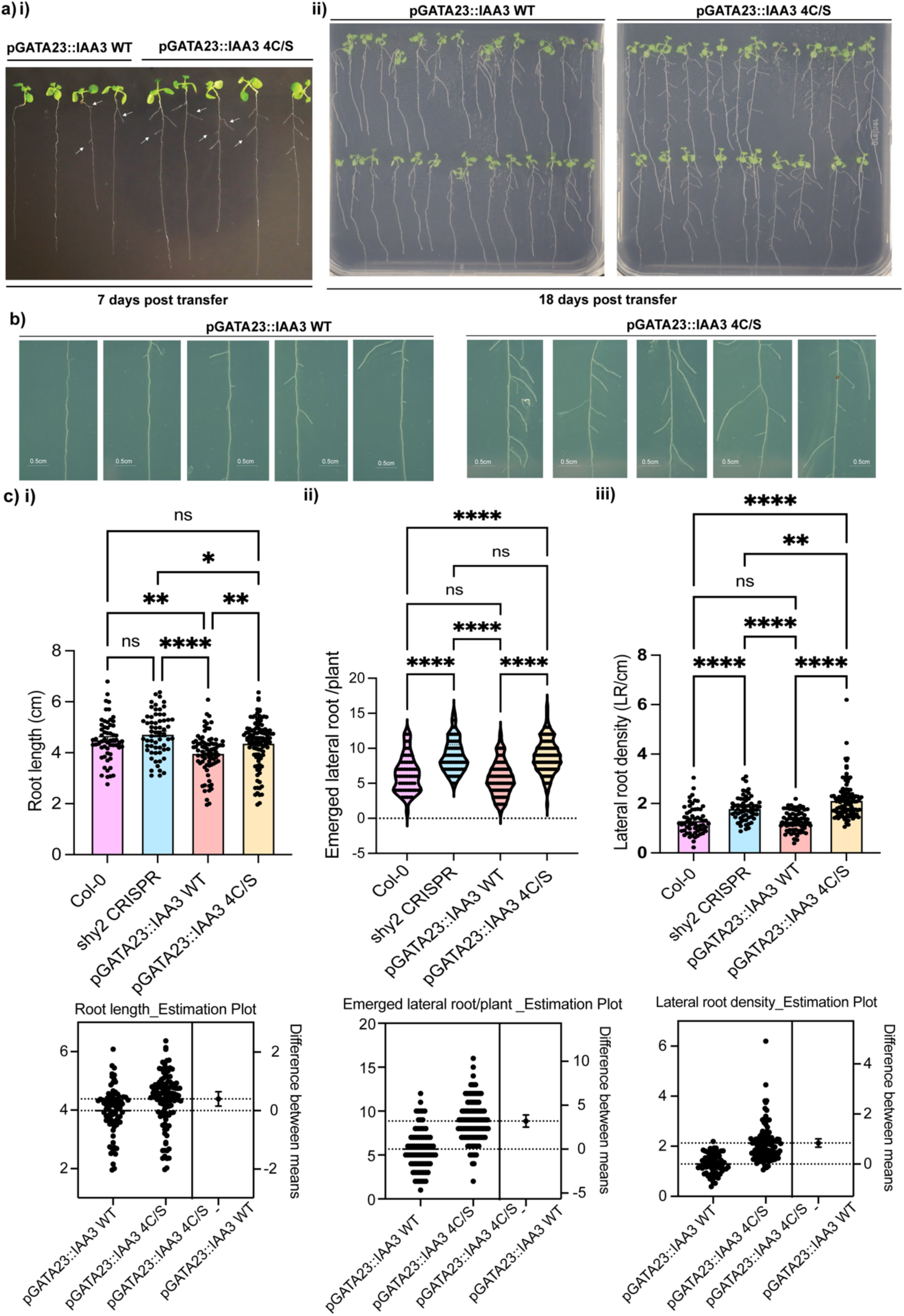
Spatio-temporal regulation of IAA3/SHY2 multimerization governs lateral root initiation Lateral Root (LR) phenotype of pGATA23::IAA3-WT and pGATA23::IAA3-4C/S expressing transgenic seedlings. (a,b) Representative images highlighting the difference in the number of emerged lateral roots per plant between pGATA23::IAA3-WT and pGATA23::IAA3-4C/S lines (c) Comparative analysis of the differences in (i) root length,(ii) number of emerged lateral roots per plant,(iii) lateral root density between WT col0, shy2 CRISPR, pGATA23::IAA3-WT and pGATA23::IAA3-4C/S lines. ns, not significant, *P ≤ 0.05; ***P≤ 0.001 ****P ≤ 0.0001.

We next tested whether quadruple cysteine mutations disrupt IAA3/SHY2 function apart from its capacity to form robust multimers in a redox dependent manner. This included the ability of 4C/S-IAA3/SHY2 to interact with ARF7^18^ and TIR1^19–22^ by transient co-immunoprecipitation experiments in *Nicotiana benthamiana*. Pull-down assays revealed that the interaction between ARF7-GFP with WT-IAA3/SHY2 and 4C/S-IAA3/SHY2 **(Figure 4a)** and TIR1-GFP with WT-IAA3/SHY2 and 4C/S-IAA3/SHY2 (**Figure 4b)** was equivalent. Hence, the quadruple cysteine mutations do not disrupt the ability of IAA3/SHY2 to interact with core ARF and TIR1 components of the nuclear auxin signalling machinery. Furthermore, we observed that the interaction of IAA3/SHY2 with ARF7 is determined by the SUMO -SIM interaction module^16^, irrespective of the multimeric state of IAA3/SHY2 **(Supplementary figure S9)**. Crucially, our pull-down assays revealed that 4C/S-IAA3/SHY2 are significantly less efficient in their interaction with the transcriptional co-repressor TOPLESS(TPL)^23^ **(Figure 4c**). Consistent with altered repressive ability, downstream targets of IAA3/SHY2^24,25^ exhibit differential expression profiles in WT-IAA3/SHY2 and 4C/S-IAA3/SHY2 transgenic lines **(Figure 4d, 4e and Supplementary Figures S10-S12**). For example, expression levels of auxin inducible *LBD* genes *(LBD16* and *LBD29*)^26,27^ was significantly higher in pCASP1::4C/S-IAA3/SHY2 versus pCASP1::WT-IAA3/SHY2 **(Figure 4d and 4e)**). We also observed significantly higher and faster inducibility of *LBD16*^27^ in the IAA3/SHY2-4C/S transgenic lines compared to IAA3/SHY2-WT on exogeneous auxin (1-NAA) treatment **(Supplementary figure S12**). The elevated inducibility of *LBD16* in the IAA3/SHY2-4C/S transgenics reveals that the repressive function of IAA3/SHY2 is impaired by the introduction of the quadruple cysteine mutation. Hence, our phenotypic, target gene expression and pull-down data reveals how ROS responsive multimerization influences the repressive function of IAA3/SHY2 (summarized in **Figure 4f** model).

**Figure 4.**
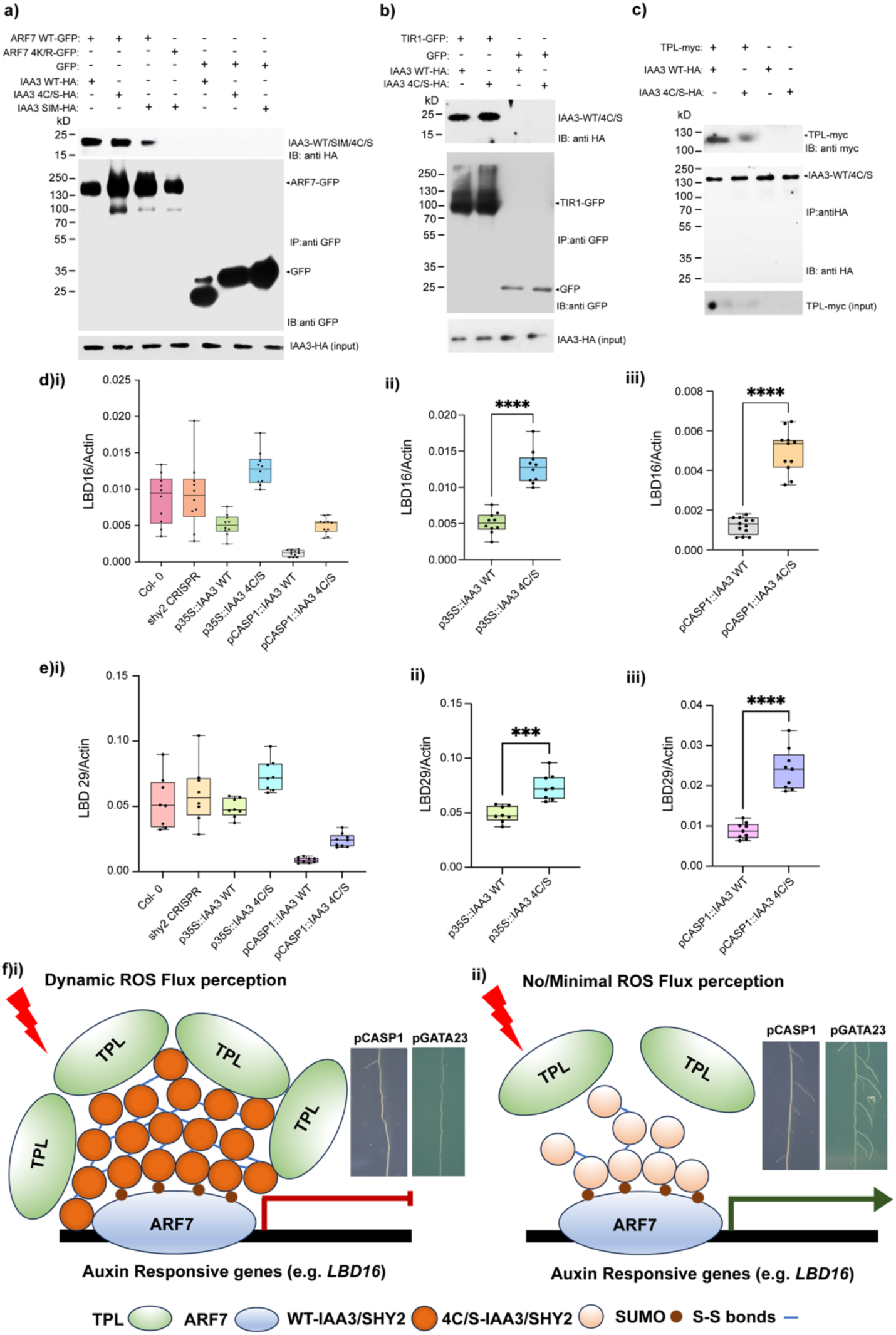
IAA3/SHY2 multimerization enhances interactions with transcriptional regulator TPL to repress downstream targetsAnalysis of the capacity of IAA3-4C/S to interact with ARF7, TIR1 and TOPLESS and mediate repression of downstream target genes. (a) Both WT and 4C/S version of IAA3 remains able to interact with ARF7-WT. Interaction of ARF7-4K/R and IAA3-SIM mutant is assessed to analyse the significance of the SUMO-SIM interaction module in mediating the interaction with ARF7. (b) IAA3-4C/S interacts with TIR1 in a manner that is similar to that of IAA3-WT indicating auxin modulated co-receptor engagement of IAA3/SHY2 is largely unaltered due to the quadruple cysteine to serine substitution mutation. (c) CO-IPs indicate that IAA3-4C/S interacts with TPL at significantly lower levels in comparison to that of IAA3-WT(d) and (e)Expression profile of representative downstream targets of SHY2/IAA3 in the absence of exogeneous auxin treatment, where (d) corresponds to LBD16 and (e)corresponds to LBD29. qRT-PCR analysis was done using the absolute quantification method and the points within the box plots indicates replicates of individual experiments done using at least two independent lines for IAA3-WT and IAA3-4C/S expressing transgenics. (f) Model of the proposed mechanism of action of ROS modulated multimerization in mediating IAA3/SHY2 repression. (i) Dynamic changes in the cellular redox potential that is generated through changes in the flux of endogenous and exogenous ROS that can be modulated with regards to the water status (homogeneous and asymmetric water distribution) is perceived by WT-IAA3/SHY2. This leads to redox dependent multimerization of WT-IAA3/SHY2 resulting in the formation of higher order multimers nucleating at target sites (e.g., *LBD16* promoter region), where they are recruited through their interaction with ARFs like ARF7, via the SUMO-SIM interaction module. Nucleation of these WT-IAA3/SHY2 higher order multimers at target sites provides an increased interface for interaction with a larger number of the TOPLESS (TPL) co-repressor molecules, leading to stronger and more robust repression. (ii) Perception of the dynamic changes in the cellular redox potential by the quadruple cysteine mutated version of IAA3/SHY2 (4C/S-IAA3/SHY2) is skewed due to the attenuation of the ability of 4C/S-IAA3/SHY2 to form higher order redox modulated multimers. Relatively weaker and less stable multimerization of 4C/S-IAA3/SHY2 thus results in significantly reduced co-recruitment of TPL co-repressor molecules at the target locus ((e.g., *LBD16* promoter region), thereby leading to de-repression of gene expression.

In summary, we report an important new layer of ROS regulation to modulate auxin-dependent transcriptional responses. We have discovered how changes in redox intersect with auxin perception by targeting selected Aux/IAAs to multimerise and repress auxin response. Our study has revealed that different Aux/IAAs act as ROS sensors with varying propensities for multimerizing through cysteine disulphide linkages dependent on their redox responsiveness. Redox dependent multimerization provides an alternate route for the Aux/IAAs to efficiently repress their downstream targets.

## Methods

### Recombinant protein purification

Fragment corresponding to the coding sequence of full-length Aux/IAAs from *Arabidopsis thaliana* and the 4C/S-IAA3/SHY2 coding sequence (generated by site directed mutagenesis of AtIAA3/SHY2 coding sequence) was cloned in pMal c5X vector (New England Biolabs) and expressed in BL21 (DE3) cells in the presence of 0.5 mM IPTG at 37 °C as an in-frame fusion with N-terminal Maltose binding protein (MBP) tag. Recombinant proteins were purified using MBP-amylose resin-based affinity chromatography under non-reducing conditions. Briefly pelleted cells were lysed with Bug buster (Merck-Millipore) protein extraction reagent (5mg Bug buster solution per gram of pellet) and the slurry was centrifuge at 5,000 rpm for 30 minutes. The supernatant was diluted one-fold with 2X column buffer (40mM Tris-HCl, 400mM NaCl, 2mM EDTA, 1mM PMSF), so that the final solution is equilibrated as a cellular lysate in Bug buster solution and 1X column buffer. The supernatant was incubated with 750ul of amylose resin that had been pre-calibrated in 1X column buffer for 3 hours. The slurry was passed through a 10ml gravity flow-based column (Fisher scientific) and the column was washed stepwise with 1X column buffer supplemented with 1mM maltose, 3mM maltose, 5mM maltose and eluted with 1X column buffer supplemented with 10mM maltose. The eluted fractions were pooled and dialysed against 20 mM Tris-HCl (pH ranging from 7.2 to 8.5, depending on the pI of the Aux/IAA protein), 150 mM NaCl, 1 mM EDTA, 5% glycerol and 1 mM PMSF. For the generation of exclusively reduced versions of full-length Aux/IAA proteins; the eluted fractions were dialysed in the same buffer containing 7.5 mM DTT. Purified recombinant proteins were analysed on an 8% SDS-PAGE gel using Coomassie Brilliant blue (CBB) staining and/or immunoblotting with anti-MBP antibody.

### Treatment of recombinant proteins with reducing and oxidizing agents

For diamide induced multimerization (DIM) assays, recombinant proteins were prepared as mentioned above in the absence of any reducing agent. Approximately 15μg (∼10uM) of recombinant IAA3-WT or IAA3-4C/S was treated with 7.5mM DTT for 30 minutes at room temperature. DTT treated protein samples were then subsequently treated with increasing concentrations of diamide ranging from 3mM to 15mM for 30 minutes at room temperature. The treated proteins were then processed in 1X Lamelli Dye without β-mercaptoethanol and analysed in an 8% SDS-PAGE gel by staining with Coomassie brilliant blue. For comparison of the differences in the multimerization capacity of the candidate Aux/IAAs, recombinant protein samples were first treated with 7.5mM DTT followed by treatment with different concentrations of NEM ranging from 1mM to 10mM and finally with a fixed concentration of 10mM diamide. In each case, the incubation was done at room temperature for 30 minutes. The treated proteins were then processed in 1X Lamelli Dye without β-mercaptoethanol and analysed in an 8% SDS-PAGE gel by staining with Coomassie brilliant blue.

### Protein sequence analyses and structure modelling

Multiple sequence alignment of IAA proteins was performed using MUSCLE^28^ in Jalview ^29^ and Mulialin^30^. IAA3 structure was modelled using Alpha fold. Three-dimensional structures were visualized with PyMOL software.

### Solution digestion

Lyophilised sample was resuspended in 23ul total volume in 5%SDS, 50mM Triethylamine ammonium bicarbonate (TEAB), pH 8.5 and acidified with 2.5ul of 27.5% phosphoric acid. 165ul of binding/wash buffer (90% methanol,100mM TEAB) was added to the sample and mix, transfer to the S-Trap column (Protifi, Fairport, NY, USA) and place in 1.7ml Eppendorf tube. This was then centrifuged at 4000 g to trap proteins for 30sec.The S-Trap column was further washed with 150μl of wash/binding buffer and centrifuged at 4,000g for 30 sec. This process was repeated 3 times in total. For digestion (either Trypsin or GluC) the column was placed in a new collection tubes and 20μl of protease solution (1ug per 10ug of protein in 50mM TEAB) and incubated overnight at 37°C. For elution, 40μl of elution buffer 1 (50mM TEAB) was added and the column centrifuged for 1 minute at 4000g, followed by 40μl of elution buffer2(0.2% Formic Acid), and centrifuged for 1 minute at 4000g, followed by the addition of 40μl of elution buffer 3 (50% Acetonitrile (ACN)) and further centrifugation at 4000g for 1 minute. The eluted peptides were lyosphilised prior to mass spectrometric analysis.

### In gel digestion

Coomassie stained band were excised from the gel and cut into 1mm^3^ pieces. The pieces were destained by adding 50% acetonitrile,50% ammonium bicarbonate solution and incubated for 10minutes at 37°C, followed further destaining with HPLC-grade water until the gel pieces were clear. The gel pieces were dehydrated by the addition of 100% ACN, for 10 mins at 37°C. The supernatant was removed, and the gel pieces were then air dried. Trypsin solution was added in 50mM ammonium bicarbonate, trypsin conc. 0.005μg/μl. The volume of trypsin solution used was sufficient to cover gel pieces. The resulting supernatant was lyophilised prior to mass spectrometric analysis.

### LC-MS/MS

All LC-MS/MS experiments were performed using a Dionex Ultimate 3000 RSLC nanoUPLC (Thermo Fisher Scientific Inc, Waltham, MA, USA) system and a Q Exactive Orbitrap mass spectrometer (Thermo Fisher Scientific Inc, Waltham, MA, USA). Separation of peptides was performed by reverse-phase chromatography at a flow rate of 300 nL/min and a Thermo Scientific reverse-phase nano Easy-spray column (Thermo Scientific PepMap C18, 2μm particle size, 100A pore size, 75μm i.d. x 50cm length). Peptides were loaded onto a pre-column (Thermo Scientific PepMap 100 C18, 5μm particle size, 100A pore size, 300μm i.d. x 5mm length) from the Ultimate 3000 autosampler with 0.1% formic acid for 3 minutes at a flow rate of 15 μL/min. After this period, the column valve was switched to allow elution of peptides from the pre-column onto the analytical column. Solvent A was water + 0.1% formic acid and solvent B was 80% acetonitrile, 20% water + 0.1% formic acid. The linear gradient employed was 2-40% B in 40 minutes. Further wash and equilibration steps gave a total run time of 60 minutes. The LC eluant was sprayed into the mass spectrometer by means of an Easy-Spray source (Thermo Fisher Scientific Inc.). All m/z values of eluting ions were measured in an Orbitrap mass analyzer, set at a resolution of 35000 and was scanned between m/z 380-1500. Data dependent scans (Top 20) were employed to automatically isolate and generate fragment ions by higher energy collisional dissociation (HCD, NCE:26%) in the HCD collision cell and measurement of the resulting fragment ions was performed in the orbitrap analyser, set at a resolution of 17500. Singly charged ions and ions with unassigned charge states were excluded from being selected for MS/MS and a dynamic exclusion window of 40 seconds was employed.

### Database searching

Post-run, text files generated from the raw data were converted to .mgf files and the files were submitted to the Mascot search algorithm (Matrix Science, London UK, version 2.7) and searched against a common contaminants database (cRAP_20190401.fasta) and the TAIR 10.0 database. Variable modifications of oxidation (M), deamidation (NQ) and phosphorylation (STY) and a fixed modification of carbamidomethyl (C) and N-ethylmaleimide (C) were applied. The peptide and fragment mass tolerances were set to 20 ppm and 0.1 Da, respectively. Scaffold (version Scaffold_5.1.1, Proteome Software Inc., Portland, OR) was used to validate MS/MS based peptide and protein identifications. Peptide identifications were accepted if they could be established at greater than 95.0% probability by the Scaffold Local FDR algorithm. Protein identifications were accepted if they could be established at greater than 99.0% probability and contained at least 2 identified peptides. Protein probabilities were assigned by the Protein Prophet algorithm^31^. Proteins that contained similar peptides and could not be differentiated based on MS/MS analysis alone were grouped to satisfy the principles of parsimony. Proteins sharing significant peptide evidence were grouped into clusters.

### Plant material and growth conditions

*Arabidopsis thaliana* ecotype, Columbia (Col-0) was used as the wild-type control for all experiments. The *shy2* CRISPR mutant lines used for this study was kindly provided by Dr. Kevin Bellande and Prof. Joop Vermeer (University of Neuchâtel). The nuclear RoGFP2orp1ROS reporter lines were kindly provided by Dr. Ulrike Bechtold and Prof.Phil Mullineaux. The *rboh* mutant lines^1^ were kindly shared by Prof. Xavier Draye. All relevant transgenic *Arabidopsis thaliana* lines expressing WT or mutated version of IAA3/SHY2 under the control of different promoters were generated by intogressing the appropriate DNA constructs (elaborated below) into the *shy2* CRISPR mutant background. *Arabidopsis thaliana* seed lines were surface sterilized using 50 % (v/v) bleach for 7 min and two 0.001 % Triton X-100 washes followed five washes with sterile water and then stratified at 4 °C for 48 h in the dark. For Lateral Root phenotype-based bioassays, seeds were germinated on media containing 1⁄2 MS (2.2 g/L) (Murashige and Skoog media, Sigma), 0.5 g/L MES, and 1 % phyto-agar at pH 5.7-5.8 range. Seedlings were grown vertically for a period of 8-10 days (depending on the experimental design) under continuous temperature 22 °C with a 16 h photoperiod (150 μmol m^−2^ s^−1^) before they were scored for their LR phenotype within a window period of 12-16 days post germination, involving root length, emerged LR number per plant and LR density as critical parameters.

### DNA constructs

All the constructs used in relation to transient assays-based studies and generation of transgenic materials were generated by utilizing combinations of Gateway cloning technology and restriction site-based cloning. ARF7WT-GFP, ARF74K/R-GFP, HA-IAA3 WT, HA-IAA3 SIM, TIR1-GFP constructs were previously generated by Orosa et al, 2018^18^. In order to generate the quadruple cysteine to serine substitution mutated version of IAA3/SHY2 (4C/S-IAA3/SHY2), site directed mutagenesis was performed using WT-IAA3/SHY2 pDTOPO clone as template. Primers used to introduce substitution mutations are listed in Supplemental table. The introduction of mutation was confirmed by sequencing. 4C/S-IAA3/SHY2 CDS in pDTOPO vector was further cloned in pEarleyGate 201 destination vector by LR reaction as an in-frame fusion with an N-terminal HA tag like HA-WT IAA3/SHY2 clone that had been previously generated by Orosa et al, 2018^18^. The 35S promoter driving the expression of HA-WT IAA3/SHY2 or HA-4C/S IAA3/SHY2 in the pEarleyGate 201 system was replaced with the CASP1 and GATA23 promoter by adopting the cloning strategy previously described by Michneiwicz et al, 2015^32^. LR reaction mediated recombinational cloning of the WT-IAA3/SHY2 or 4C/S-IAA3/SHY2 CDS in the pEarleyGate 201 vector system is accompanied with the excision of the chloramphenicol resistance (Cm^R^) gene and this renders the EcoRI and XhoI restriction sites flanking the CAMV 35S promoter as unique in the LR positive clones. The CASP1 and GAT23 promoter was then cloned into these unique EcoRI and XhoI sites by conventional restriction enzyme based cloning methods (as both the CASP1 and the GATA23 promoters lack EcoRI and XhoI sites) to generate a chimera with the aforementioned promoters driving the expression of the WT-IAA3/SHY2 or 4C/S-IAA3/SHY2 CDS as an in frame fusion with an N-terminal HA tag followed by an OCS (Octopine synthase) terminator.

### Generating transgenic materials

DNA constructs expressing N-terminal HA tagged WT-IAA3/SHY2 and 4C/S-IAA3/SHY2 under the control of the CAMV 35S, CASP1 and GATA23 promoter as described above were transformed into *Agrobacterium tumefaciens* GV3101 strain by electroporation. The expression of the chimeric proteins (HA-WT IAA3/SHY2 and HA-4C/S IAA3/SHY2) were first confirmed by agrobacterium mediated transient assays followed with anti -HA immunoprecipitation (described below) and then transformed into *Arabidopsis thaliana* Col-0 plants by the floral dip method^33^. Positive transformants were selected in 20mg/litre BASTA selection media (the pEarleyGate 201 destination vector contained the biolophos/ phosphonithricin resistance gene for *in planta* selection).

### Western blot analysis of transgenic plants

Total protein was isolated from Arabidopsis thaliana transgenic seedlings expressing N-terminal HA tagged WT IAA3/SHY2 and 4C/S-IAA3/SHY2 in RIPA buffer (20 mM Tris-HCl (pH 7.5), 150 mM NaCl, 1 mM Na_2_EDTA, 1% NP-40, 1% sodium deoxycholate, 2.5 mM sodium pyrophosphate, 1 mM β-glycerophosphate, 1 mM PMSF and 1 mM β-Mercaptoethanol) maintaining a tissue: buffer ratio of 1:1. Crude protein extracts were processed with 4X Lamelli Dye supplemented with β-Mercaptoethanol and separated by 10% SDS-PAGE gels and immunoblotted with the anti-HA antibody (anti-HA Abcam). For endogenous controls, immunoblotting was performed with Anti-histone H3 antibody (Abcam).

### Agrobacterium mediated transient assays

Gene constructs were transiently expressed in *Nicotiana benthamiana* plants using the *Agrobacterium tumefaciens* mediated transformation^34^. ARF7-GFP, ARF7–GFP (4K/R), TIR1-GFP and GFP proteins were used to check the interactions with HA-WT IAA3/SHY2, HA-4C/S IAA3/SHY2 and HA-SIM IAA3/SHY2 in *Nicotiana benthamiana* leaves.3 d post-infiltration, leaves were frozen in liquid nitrogen for sample preparation and co-immunoprecipitation, or immunoprecipitation assays were performed.

### Co-immunoprecipitation assay

*Nicotiana Benthamiana* plants were infiltrated with a combination of *Agrobacterium tumefaciens* cells expressing the proteins of interest along with the anti-silencing factor P19. It was ensured that the total O.D. for the combination of the *Agrobacterium tumefaciens* cells did not exceed 0.6 in all the co-immunoprecipitation (co-IP) assays. 3 days post infiltration, total protein was then extracted from *Nicotiana Benthamiana* leaves for co-IP using the extraction buffer containing: 50 mM Tris HCl (pH 8.0), 150mM NaCl, 1mM EDTA, 0.5% Trition-X 100, 2% [v/v] Glycerol and 10 mM DTT (depending on the type of immunoprecipitation performed). Anti-GFP IP and anti-HA IP were performed. Total protein was incubated with either 20 μl anti-GFP or 20 μl anti-HA micro magnetic beads (Miltenyi Biotech) and put on a rotating platform at 4°C for 30 minutes and the slurry was subsequently passed through a micro column (Miltenyi Biotech) attached to a magnetic stand (Miltenyi Biotech) to capture the beads. The beads were then washed four times by passing 1 ml of cold IP buffer through the micro column. After the last wash, 80 μl of pre-heated (98°C) 1× Lamelli buffer was used to elute the immuno-complexes. The immuno-complexes was then analysed on a 10% SDS-PAGE gel using immunoblotting methods with Abcam (Cambridge, UK) anti-GFP, anti-HA antibodies (Anti-HA High Affinity Roche 3F/10, Sigma-Aldrich, and anti-HA Abcam). At least three independent biological replicates were performed per experiment.

### Redox dependent Immunoprecipitation (Redox IP) assays

Transgenic *Arabidopsis thaliana* seedlings or leaves from *Nicotiana Benthamiana* plants infiltrated with a combination of *Agrobacterium tumefaciens* cells expressing the proteins of interest along with the anti-silencing factor P19 were used for Redox dependent Immunoprecipitation assays. Total protein from samples were isolated for Redox-IP using extraction buffer containing: 25 mM Tris-HCl (pH 8.0), 150 mM NaCl, 1mM EDTA, 0.25% [w/v] Sodium deoxycholate, 2% [v/v] glycerol, 0.1% [v/v] TritonX-100, 0.7% [v/v] Igepal CA-630 (Nonidet P-40), 50 mM KCl, 20mM MgCl_2_ and protease inhibitor cocktail. Depending on the redox state of the extraction buffer intended for the assay, the extraction buffer was supplemented either with 10mM DTT or 10mM Diamide. Anti-HA IP was then performed. Total protein was incubated with 20 μl anti-HA micro magnetic beads (Miltenyi Biotech) and put on a rotating platform at 4°C for 30 minutes and the slurry was subsequently passed through a micro column (Miltenyi Biotech) attached to a magnetic stand (Miltenyi Biotech) to capture the beads. The beads were then washed four times by passing 1 ml of cold Redox-IP buffer through the micro column. After the last wash, immunocomplexes were eluted with 80 μl of pre-heated (98°C) 1× Lamelli buffer that either lacked β-Mercaptoethanol (for redox IPs performed in non-reducing or oxidizing environment) or were supplemented with β-Mercaptoethanol (for redox IPs performed in reducing environment). The immuno-complexes was then analysed on a 10% SDS-PAGE gel using immunoblotting methods with anti-HA antibodies (Anti-HA High Affinity Roche 3F/10, Sigma-Aldrich, and anti-HA Abcam). At least three independent biological replicates were performed per experiment. For DTT based titration assays, total protein from samples were extracted in the extraction buffer described above and anti HA IP was performed. Elution of the immunocomplexes were eluted with 80 μl of pre-heated (98°C) 1× Lamelli buffer that lacked β-Mercaptoethanol, and the immuno-complexes were analysed by the same methodology as described above.

### Detection of SUMO conjugation of targets

p35S::YFP-ARF7 WT and p35S::YFP-ARF7^4K/R^ constructs were transformed into *Agrobacterium* strain GV3101 and were coinfiltrated along with p35S::HA-SUMO1 and P19 into *N.benthamiana*. The infiltrated *N.benthamiana* leaf samples were collected after 3 days, and total protein was isolated from the samples in extraction buffer containing: 50 mM Tris-HCl (pH 8.5), 150 mM NaCl, 1mM EDTA, 0.1% [v/v] SDS, 0.5% [w/v] Sodium deoxycholate, 1.0 % [v/v] Igepal CA-630 (Nonidet P40), 50mM KCl, 50mM NEM (N-ethyl maleimide) and protease inhibitor cocktail. Anti-GFP IP was performed with the total protein extract using the methodology as described above, and the immuno-complexes was then analysed on a 10% SDS-PAGE gel using immunoblotting methods with Abcam (Cambridge, UK) anti-GFP and anti-HA antibodies (Anti-HA High Affinity Roche 3F/10, Sigma-Aldrich, and anti-HA Abcam). At least three independent biological replicates were performed per experiment.

### Agar-based xerobranching assay

Agar-based Air-gap Assay was designed for phenotyping *Arabidopsis* thaliana seedlings exposed to transient water stress in accordance with Mehra et al, 2022^3^. Briefly, ½ MS medium supplemented with 0.3 % sucrose, 0.5 g/L MES and 1 % Bacto agar was poured into square petri plates (12 cm x 12 cm), and agar slants were created by aligning Petri dishes at a 15° angle with respect to a flat surface. In the middle of the petri dish, a clean cut was made with the help of a 10-cm long scraper blade, and cut agar was carefully removed using a 0.5 scalpel, to create an air-gap of ∼0.5 cm. Five days-old *Arabidopsis* seedlings were placed on plates in a manner that allowed the root tips to be positioned 2-3 mm away from the agar-air interface. Plates were sealed and held vertically under appropriate growth conditions (as described above) for 7 days.

### Statistical Analysis

Both Student’s *t*-test and ANOVA was used to perform the statistical analysis of the data. The Student’s *t-*test determined if the means of two data sets differed significantly from each other in relation to the null hypothesis. A different letter indicated a significant difference from that of WT (Col-0) roots based on the Student *t*-test *P* value (*P*<0.05). ANOVA was used when there were more than two data sets to compare. The analysis of variance, one-way ANOVA considered the total number of observations, means, standard deviations (or standard error) within each data set. Using GenStat, multiple comparisons, Tukey HSD (Honest Significant Difference) test was applied at the *P*-value 0.05 significance level (95% confidence interval) to each data set. Asterisk notation was used to indicate significance levels between all the data sets in relation of analysis of LR phenotype and quantitative real time PCR data. Additionally, for analysis of parameters related to LR phenotype (root length, emerged lateral root per plant and lateral root density) in normal ambient water supplied conditions, Welch’s unequal variance t-test was also performed to consider unequal population variances in a two-sample location test. Transgenic seedlings expressing WT-IAA3/SHY2 and 4C/S-IAA3/SHY2 under the control of different promoters were used as the two different populations to test the null hypothesis that these populations have equal means. Statistical analysis was done using software Prism-GraphPad. In relation to the data corresponding to Xerobranching assays, letters were used to indicate significance levels between all the data sets, where similar letters stated that there were no significant differences observed. Whereas different letters indicated that there was a significant difference observed between the data sets (*P*<0.05).

### Confocal imaging and image analysis

Imaging of roGFP2-ORP1 was performed as described previously (Nietzel et al., 2018 New Phytologist). Roots exposed to control conditions, XB stimuli and recovery conditions were imaged using a Leica SP5 confocal laser scanning microscope (Leica Microsystems). The GFP signal was sequentially excited at 402 nm and 488 nm wavelengths and emission was recorded at 505-535 nm. Images were acquired as Z-stacks in 12-bit mode. Images were processed to calculate 402/488 ratios using ImageJ (ImageJ 1.53f51).

### RNA isolation and Quantitative real time PCR (q-RT PCR)

Total RNA was extracted from 12 to 14-day old seedlings grown in 1⁄2 MS media grown vertically for a period of 8-10 days (depending on the experimental design) under continuous temperature 22 °C with a 16 h photoperiod (150 μmol m^−2^ s^−1^). A Direct-zol RNA Miniprep kit (ZYMO research) was used to extract total RNA following the manufacturer’s recommendations. Cross contaminating DNA was removed from the samples by on-column DNase I digestion following the manufacturer’s recommendations. The RNA was then quantified by measuring the absorbance (260 and 280 nm) using a NanoDropTM 1000 Spectrophotometer (Thermo Scientific), and cDNA was synthesized with Invitrogen SuperScript® II Reverse Transcriptase. The synthesized cDNA was subsequently used as template for quantitative real time PCR based analysis of the expression levels of the target genes using Brilliant III Ultra-Fast SYBR QPCR MM (Agilent) in conjunction with Rotor-Gene® Q (Qiagen) system. In order to assess the expression levels of the transgene (HA-IAA3 / SHY2 CDS expressed under the control of different promoters) and the auxin responsive downstream targets of IAA3/SHY2 in the transgenic lines generated in this study, we adopted the absolute quantification-based approach for our quantitative real time PCR based analysis in accordance with Paul et al, 2017^35^. Briefly, forward and reverse primers corresponding to the HA tag and 3’ end of the IAA3/SHY2 CDS was used to uniquely amplify the transgene (HA-WT IAA3/SHY2 or HA-4C/S IAA3/SHY2) intogressed in the *shy2* CRISPR background and expressed under the control of the CAMV 35S, CASP1 and GATA23 promoter. Logarithmic fold dilutions of plasmids cloned with HA-WT IAA3/SHY2, HA-4C/S IAA3/SHY2 CDS and an amplicon corresponding to the CDS of the housekeeping gene ACTIN7 (At5g09810) were used as template for real time PCR reaction and Ct values were plotted against the dilutions to generate a standard curve for both IAA3/SHY2 CDS and actin. The levels of the transgene in the different transgenic lines were then calculated by extrapolation of the Ct value against the generated standard curve followed by normalisation with the actin levels. WT col0 and *shy2* CRISPR seedlings were used as negative controls for the analysis. Based on this approach, transgenic lines showing similar levels of HA-WT IAA3/SHY2 and HA-4C/S IAA3/SHY2 expression at the transcriptional level were selected and the abundance of the HA-WT IAA3/SHY2 and HA-4C/S IAA3/SHY2 proteins were further verified. Post verification, multiple optimised transgenic lines were used to assess the expression levels of the auxin responsive downstream targets of IAA3/SHY2 in the absence of exogeneous auxin treatment, using the same approach as described above with the exception that the standard curves for each target gene tested was generated from gel extracted DNA fragments corresponding to the amplicon of the target gene. To assess difference in the inducibility of LBD16 expression on exogeneous auxin treatment between the transgenic lines, conventional ΔΔCt method^36^ was adopted with the housekeeping gene ACTIN7 (At5g09810) used for normalisation. Fold change was calculated relative to untreated control.

### Lateral root density

Images of whole plates were taken using a Canon digital camera. PR length was measured using the NeuronJ plugin in ImageJ. LR number was counted manually and LR density was calculated by dividing LR number by the length of the root.

## ACKNOWLEDGMENTS

We thank P. Mullineaux (University of Essex) and U. Bechtold (University of Essex and University of Durham) for kindly providing the RoGFP2 orp1 fluorescent reporter mutants. We thank Prof. Xavier Draye for kindly providing the *rboh* mutants.

## FUNDING

This study was supported by Biotechnology and Biological Sciences Research Council (BBSRC) grant (BB/T001437/1 to D.R.) and (BB/V003534/1 to D.R.); European Molecular Biology Organization Long-Term Fellowship (ALTF 1140-2019 to P.M.); European Union’s Horizon 2020 research and innovation program, Marie Skłodowska-Curie, XEROBRANCHING (891262 to P.M.), Global Challenge Research Fund (GCRF) fellowship (UKRI GCRF Durham PhD scholarship award to V.M.); Biotechnology and Biological Sciences Research Council (BBSRC) grant (BB/V003534/1 to L.C.); the Biotechnology and Biological Sciences Research Council grant (BB/T001437/1 to M.J.B. and A.S).; (BB/V003534/1 to M.J.B. and A.S).; (BB/W008874/1 and BB/W015080/1 to M.J.B);European Research Council grant FUTUREROOTS (294729 to M.J.B.); European Research Council, SUMOrice (grant no. 310235 to A.S.); ChromAuxi grant from the Agence Nationale de la Recherche (ANR-18-CE12-0014-02 to T.V. and R.M.A); Biotechnology and Biological Sciences Research Council (BBSRC) grant (BB/V003534/1 to K.S. and A.J.); Biotechnology and Biological Sciences Research Council (BBSRC) grant (BB/V003534/1 to K.L).

## AUTHOR CONTRIBUTIONS

Conceptualization: D.R., A.S., and M.J.B. Methodology: D.R., P.M., V.M., L.C., R.M.A., K.S. Investigation: D.R., P.M., V.M., L.C., K.B., R.M.A, K.S., A.B. Funding acquisition: A.S. and M.J.B. Supervision: A.S. and M.J.B. Writing – original draft: D.R., A.S. and M.J.B. Writing – review & editing: D.R., P.M., V.M., L.C., K.B., J.V., R.M.A, T.V., K.S., A.J., U.B., P.M., K.L., A.B., M.J.B and A.S.

## Competing interests

The authors declare no competing interests.

## Data and materials availability

No restrictions are placed on materials, such as materials transfer agreements. All data are available in the main text or the supplementary materials.

## Additional Information

Materials and Methods Supplementary Figures. S1 to S13 Tables S1 to S2

Excel sheet 1: List of interactors of WT -IAA3/SHY2 and 4C/S-IAA3/SHY2 (pCASP1 promoter)

## Extended data figure/table legends

**Supplementary Figure S1.**
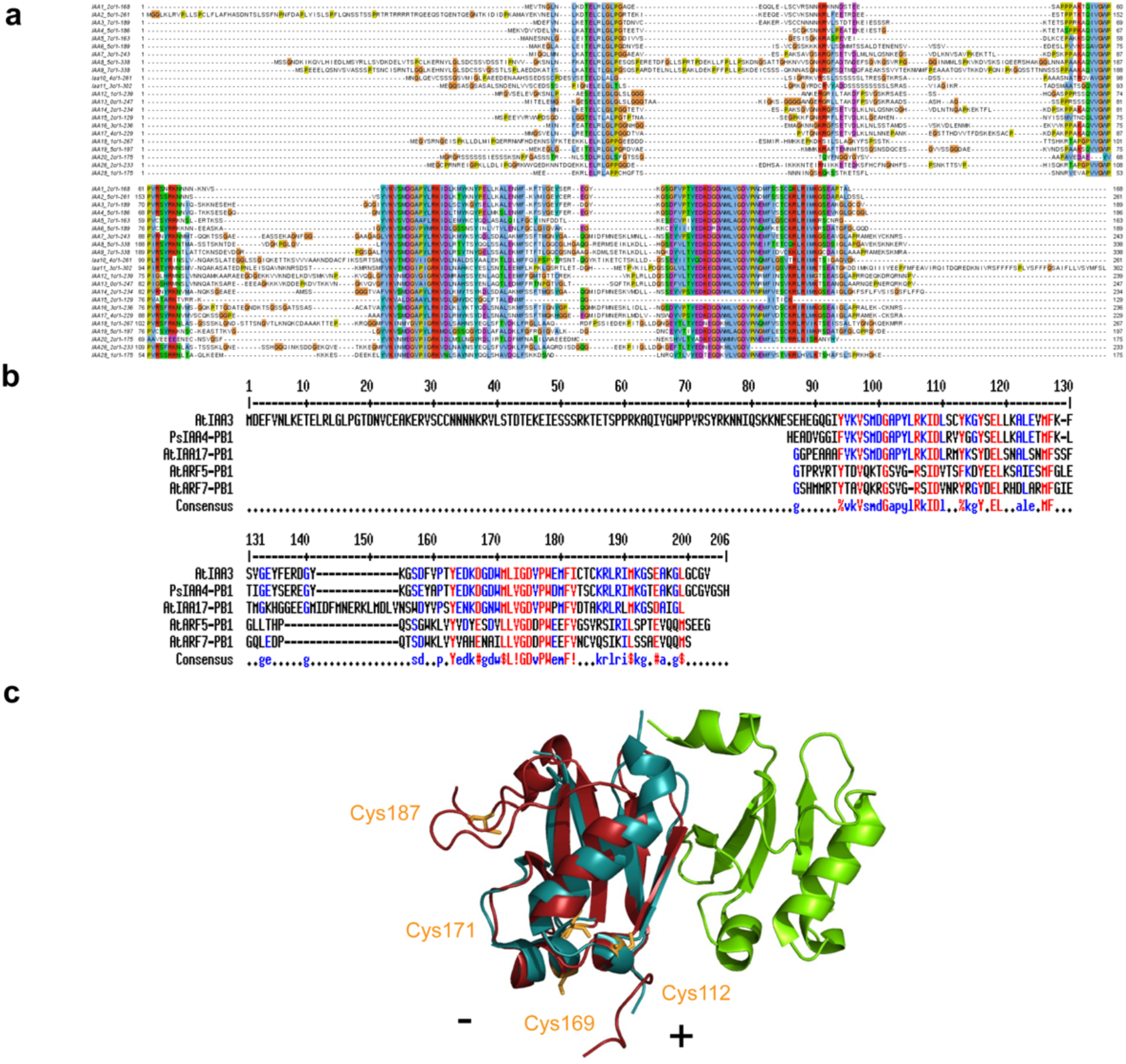
In silico analysis predicting the cysteine residues of IAA3/SHY2 involved in multimerization. **(a)** Sequence alignment of Aux/IAAs highlighting the conservation of the cysteine residues. Cysteine residue 171 is the only conserved residue **(b)** Alignment of Aux/IAAs and ARFs with known crystal structures of their PB1 domain **(c)** Modelled structure of IAA3 PB1 domain (red) super imposed on the crystal structure of IAA17 (cyan) -ARF5 (green) complex PB1 domain^37^ *(pdb 6L5K, reference: Determinants of PB1 Domain Interactions in Auxin Response Factor ARF5 and Repressor IAA17 Kim et al., 2020).* Position of the cysteine residues of IAA3/SHY2 predicted to be involved in the disulphide linkage mediated multimerization is highlighted in orange. “-” and “+” signs indicate the negative and positive interaction surfaces of the PB1 domain.

**Supplementary Figure S2.**
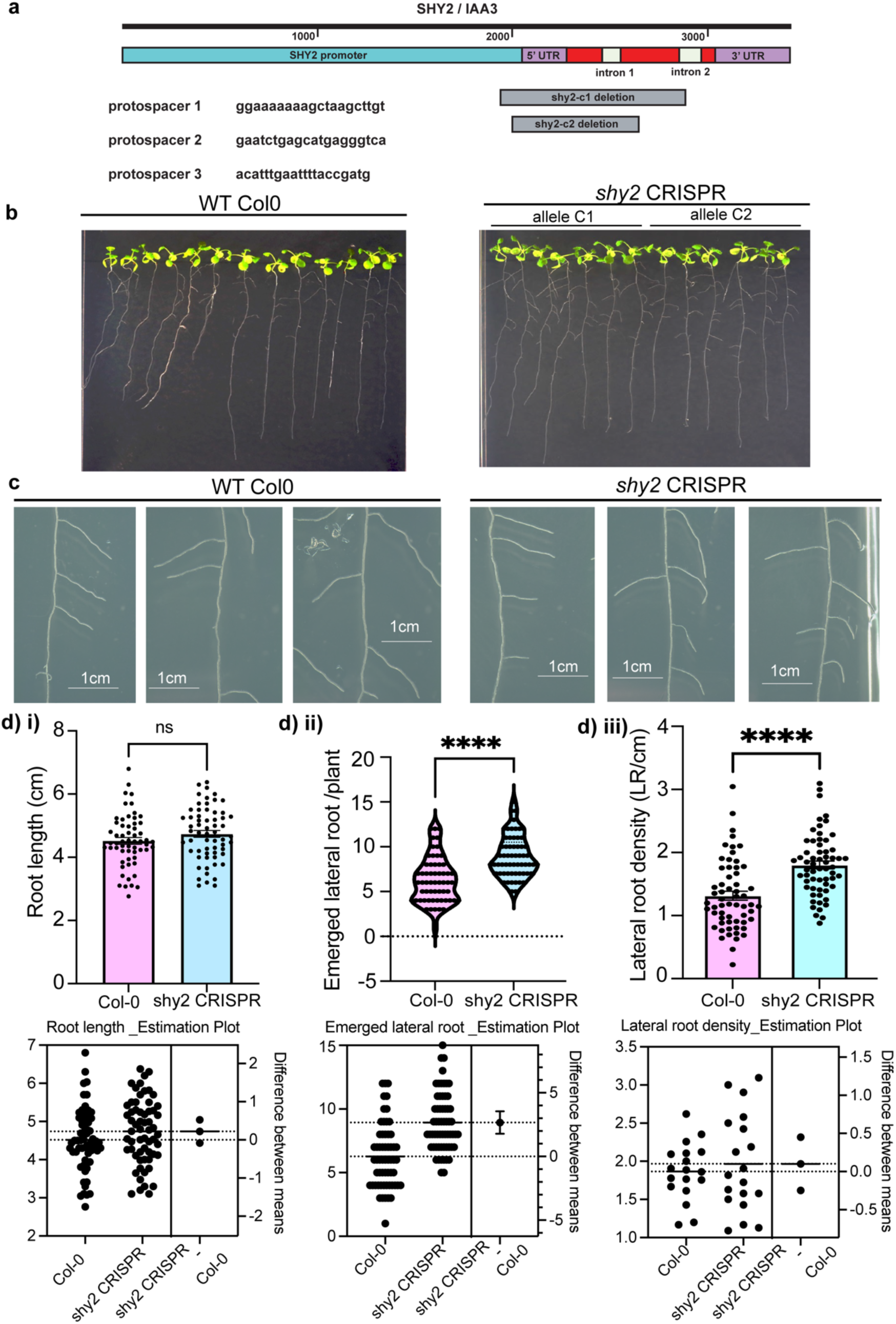
s*hy2* CRISPR deletion mutants are affected in lines have a lateral root developmentphenotype. **(a)** Schematic representing CRISPR cas9 mediated deletion at the SHY2 locus **(b, c)** shy2 CRISPR lines have higher emerged lateral roots per plant in comparison to WT Col0 **(d)** Comparative analysis of the differences in (i) root length, (ii) number of emerged lateral roots per plant, (iii) lateral root density between WT col0 and shy2 CRISPR lines. ns, not significant, *P ≤ 0.05; ****P ≤ 0.001; ns, nonsignificant changes.

**Supplementary Figure S3:**
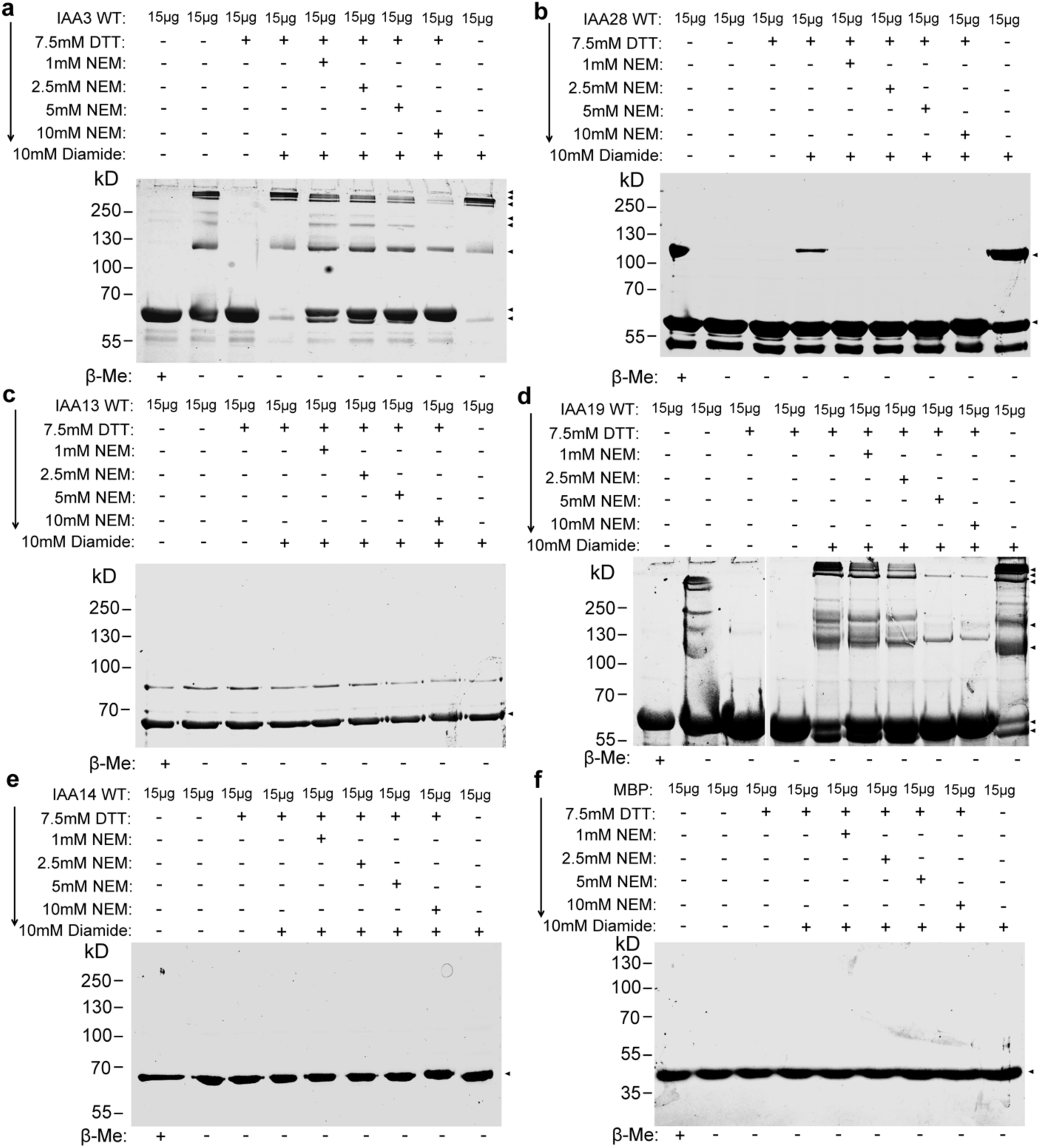
Analysis of the sensitivities of different Aux/IAAs to redox changes. Multimerization profile of **(a)** IAA3 **(b)** IAA28 **(c)** IAA13 **(d)** IAA19 **(e)** IAA14 and **(f)** MBP. Full length native protein was treated with 7.5mM DTT alone (lane3), or with 10mM Diamide after treatment of the full-length protein with 7.5mM DTT (lane4), or with10-mM diamide after treatment of the full-length protein first with 7.5mM DTT, followed by treatment with gradually increasing concentrations of NEM (N-ethyl maleimide) ranging from 1mM to 10mM NEM (lane5-8). Treatment of the different full-length Aux/IAAs with 10mM diamide alone (lane 9) is used to assess the sensitivity of the Aux/IAAs to an exclusively oxidizing environment. Electrophoresis of the treated protein samples were done in SDS PAGE gels in the absence of β-Me except for the samples loaded in lane 1 as experimental control. Direction of the arrowhead indicates the sequence in which the redox reagents are added in the assay. Aux/IAAs exhibiting higher order multimers even at higher threshold concentrations of NEM indicate that these Aux/IAAs have a higher frequency of cysteine residues that are likely to be engaged in disulphide linkages.

**Supplementary Figure S4:**
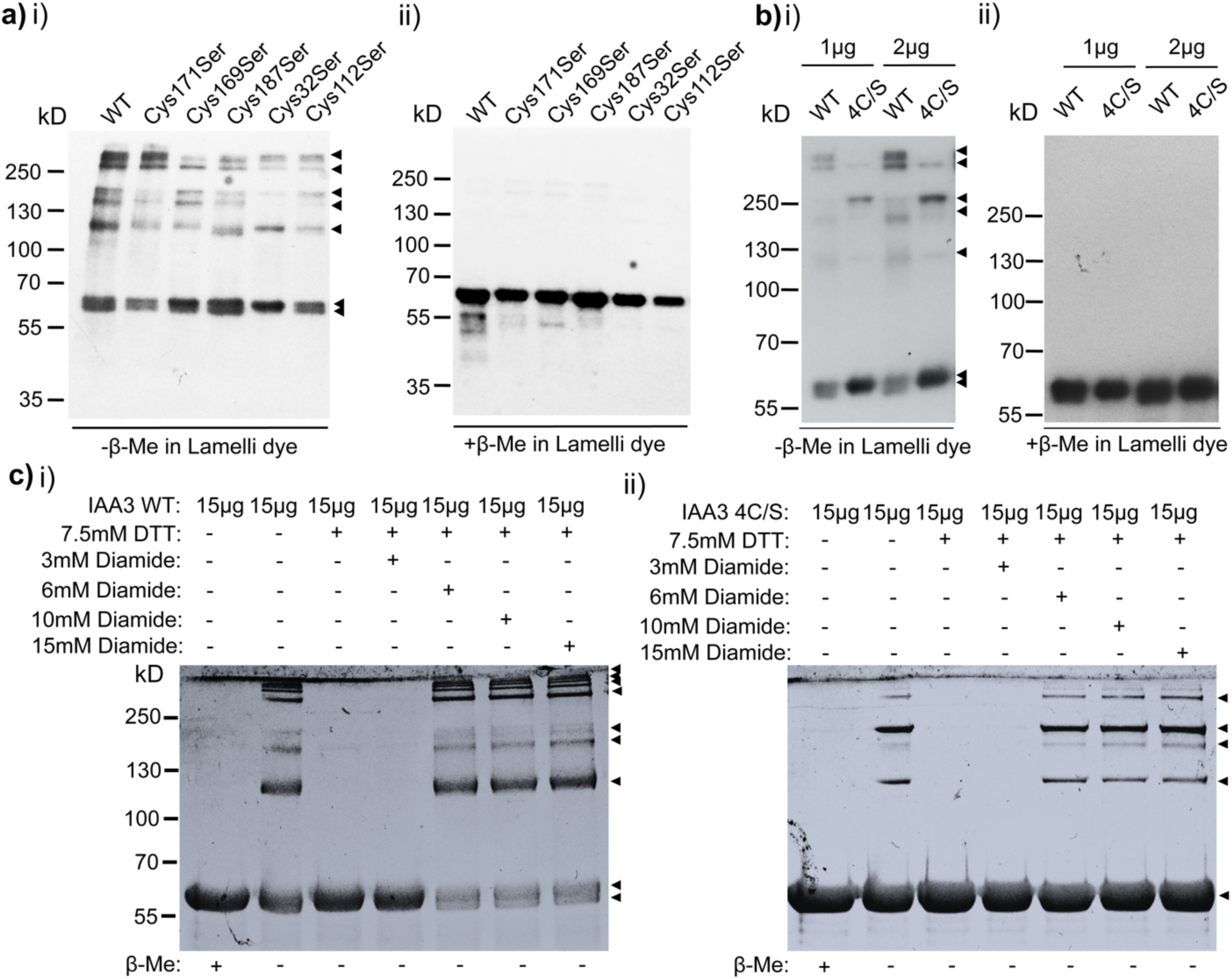
Identification of the key cysteine residues responsible for redox dependent IAA3 multimerization. **(a)** Impact on the multimerization of IAA3 due to single cysteine to serine substitutions **(b)** Impact on the multimerization of IAA3 due to combinatorial cysteine to serine substitutions (IAA3-4C/S) **(c)** Reoxidation of DTT reduced IAA3-4C/S by the thiol specific oxidizing agent, diamide, results in an alternate pattern of multimerization compared to IAA3 WT. Major fraction of IAA3-4C/S is monomeric even in the presence of 15mM diamide. + or - β-me indicates the presence or absence of β-mercaptoethanol in Lamelli dye.

**Supplementary Figure S5.**
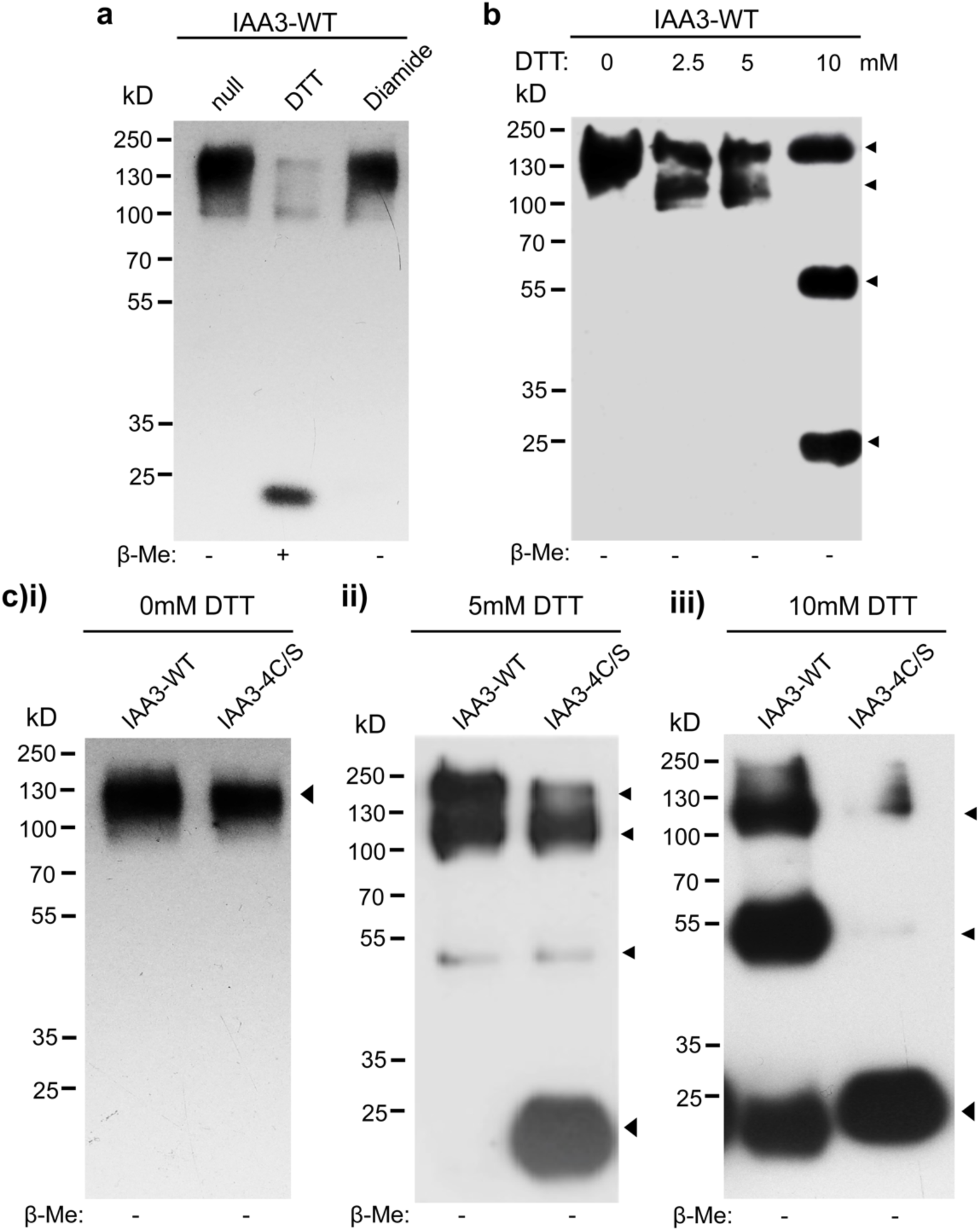
Redox dependent multimerization analysis of IAA3-WT and IAA3-4C/S in *Nicotiana benthamiana*. **(a)** Multimerization profile of IAA3 WT in buffer systems that are non-reducing (null), reducing (DTT) or oxidizing (diamide) **(b)** Multimerization profile of IAA3 WT in buffer systems with increasing concentration of DTT (upto 10mM). **(c)** Comparative analysis of the robustness of multimerization between IAA3-WT and IAA3-4C/S using buffer systems having increasing amounts of DTT. + or - β-Me indicates the presence or absence of β-mercaptoethanol in Lamelli dye used for elution and electrophoresis. All the immunoprecipitation was done with anti-HA microbeads and the immunoblotting was done with anti-HA antibody (Roche3F/10).

**Supplementary Figure S6.**
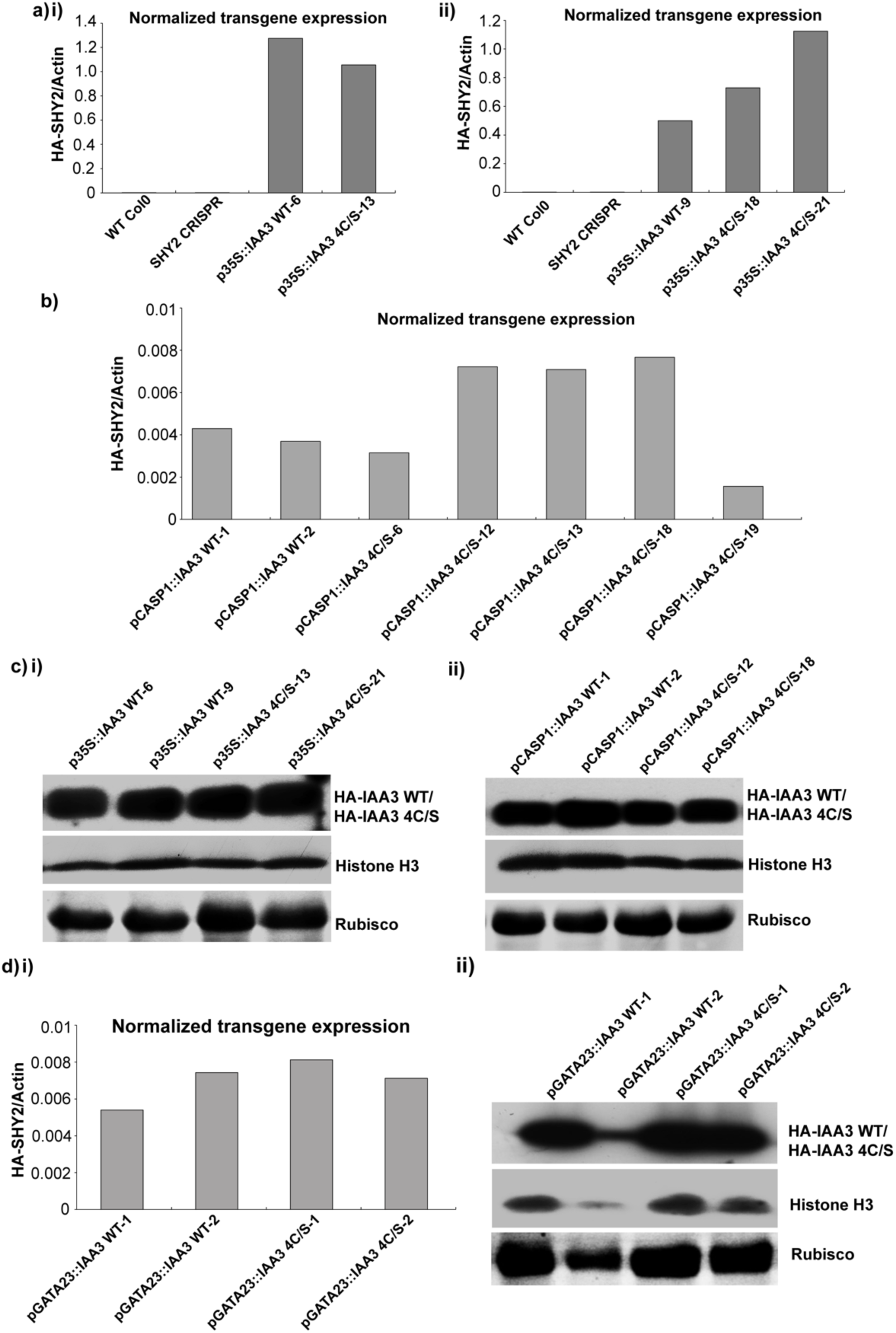
Expression analysis of the transcript and protein levels of HA-tagged WT SHY2/IAA3 and 4C/S mutant transgenic lines Assessment of the expression of the transgene HA-SHY2 at the transcriptional level. **(a, b)** and at the translational level **(c)**. Lines having comparatively equal protein abundance were taken forward for subsequent analysis. The y axis scale for graphs in **(a, b)** are universal and reflects the difference in the expression of the transgene under the control of different promoters. Gene expression levels were assessed by the absolute quantification method with a plasmid harbouring the CDS of SHY2 in fusion with the DNA sequence of the HA-tag used to generate the standard curves. The abundance of the protein levels of HA-IAA3-WT and HA-IAA3-4C/S in **(c)** was assessed by probing protein extracts from transgenic seedlings with anti-HA antibody (Roche 3F/10). Histone H3 and Rubisco were used as loading controls. Wt Col0 and SHY2-CRISPR were used as negative reference controls.

**Supplementary Figure S7.**
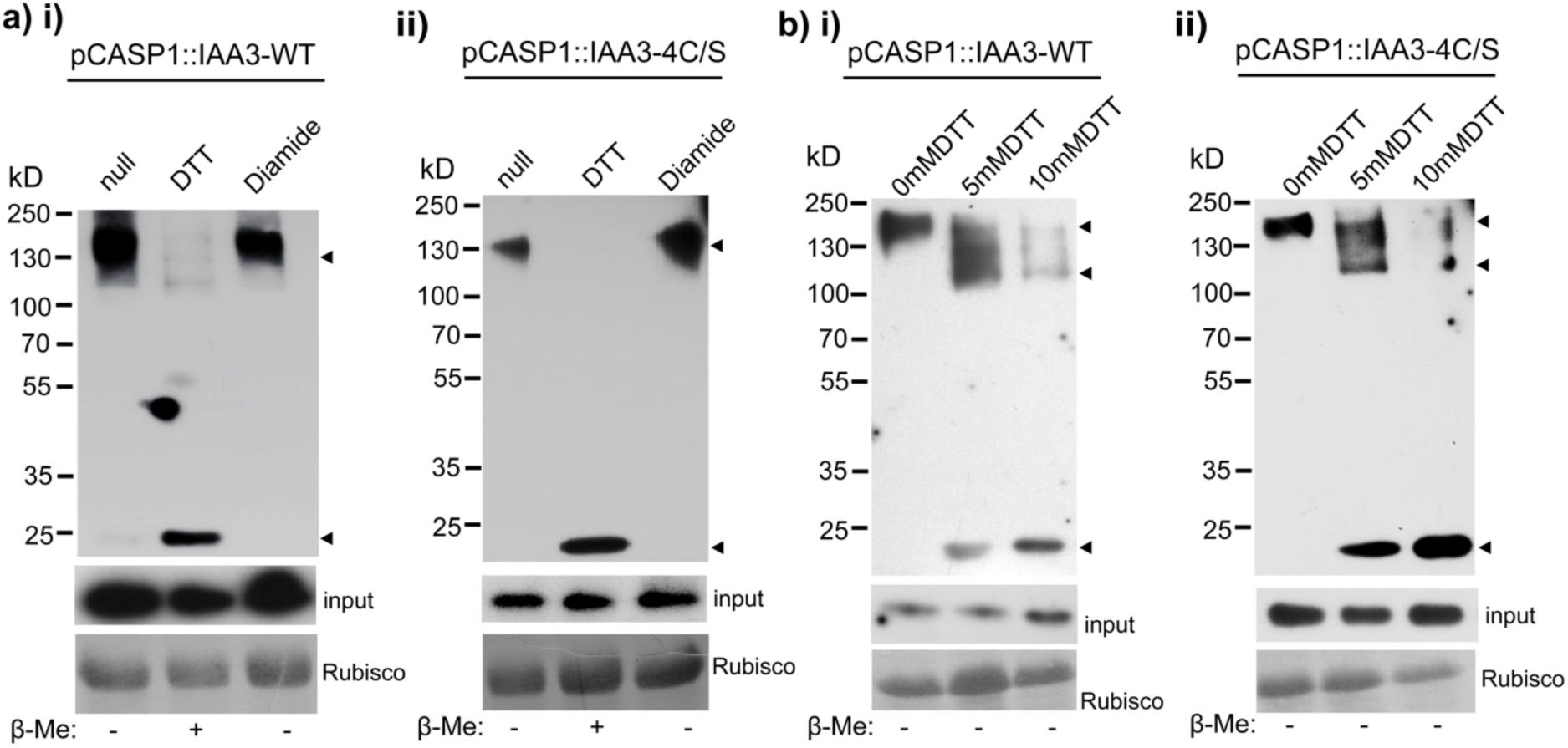
Redox dependent multimerization analysis of IAA3-WT and IAA3-4C/S expressed under the control of the CASP1 promoter. **(a (i) and (ii))** Multimerization profile of IAA3 WT and IAA3-4C/S expressed under the control of the CASP1 promoter in transgenic Arabidopsis thaliana seedlings using buffer systems that are non-reducing (null), reducing (DTT) or oxidizing (diamide) **(b (i) and (ii))** Comparative analysis of the robustness of multimerization between IAA3-WT and IAA3-4C/S using buffer systems having increasing amounts of DTT.+ or - β-Me indicates the presence or absence of β-mercaptoethanol in Lamelli dye used for elution and electrophoresis. All the immunoprecipitation was done with anti HA microbeads and the immunoblotting was done with anti HA antibody (Roche 3F/10).

**Supplementary figure S8.**
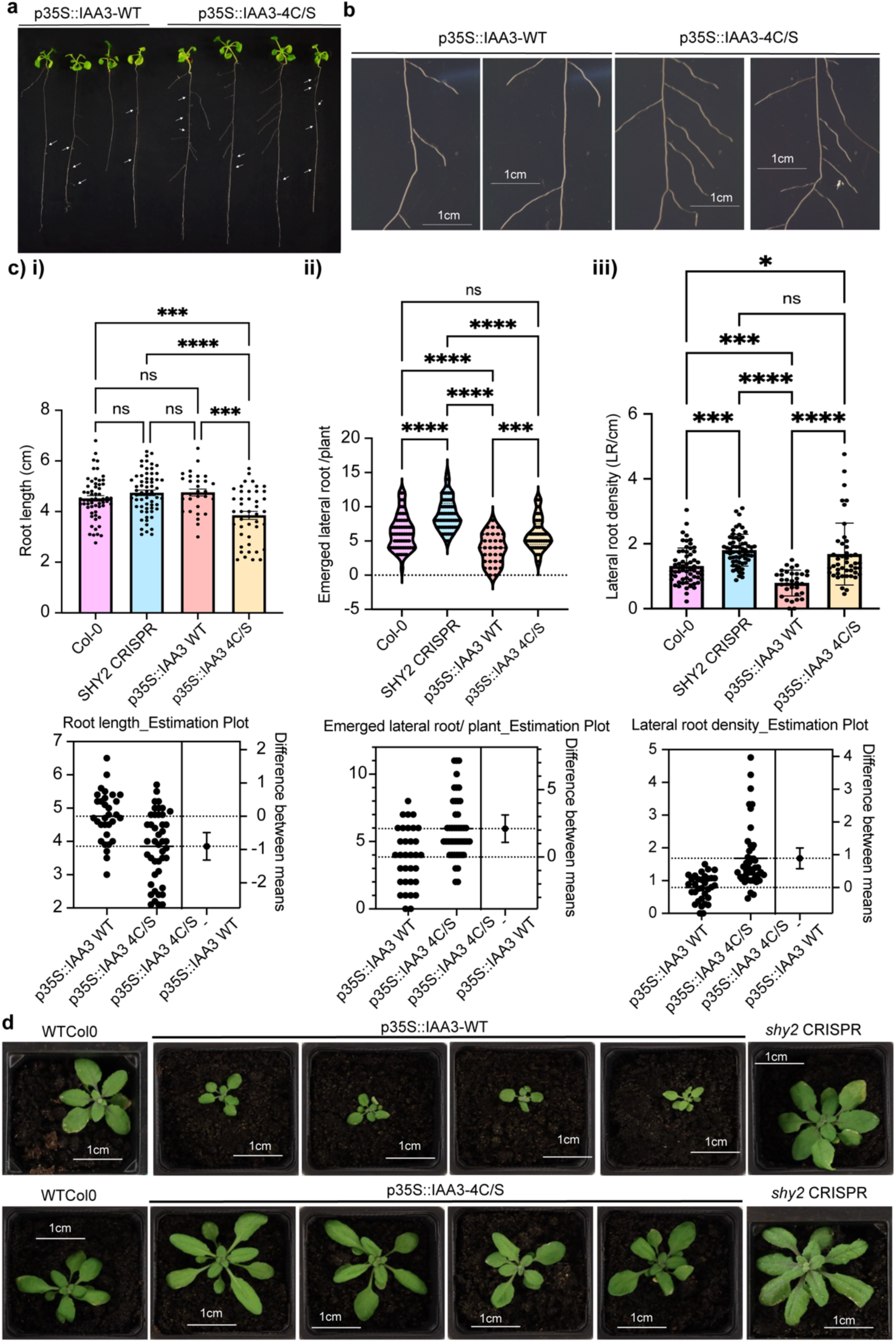
Lateral Root (LR) and rosette size phenotype analysis of shy2 CRISPR mutants expressing IAA3 or 4C/S mutant driven by the 35S promoter. Lateral Root (LR) and rosette size phenotype of p35S::IAA3-WT and p35S::IAA3-4C/S expressing transgenic seedlings. **(a,b)** Representative images highlighting the difference in the number of emerged lateral roots per plant between p35S:: IAA3-WTand p35S::IAA3-4C/S lines **(c)** Comparative analysis of the differences in (i) root length,(ii) number of emerged lateral roots per plant,(iii) lateral root density between WT col0, shy2 CRISPR, pCASP1::IAA3-WT and pCASP1::IAA3-4C/S lines. ns, not significant, *P ≤ 0.05;***P≤ 0.001 ****P ≤ 0.0001; **(d)**Representative images highlighting the difference in the rosette diameter between p35S::IAA3-WTand p35S::IAA3-4C/S lines.

**Supplementary Figure S9.**
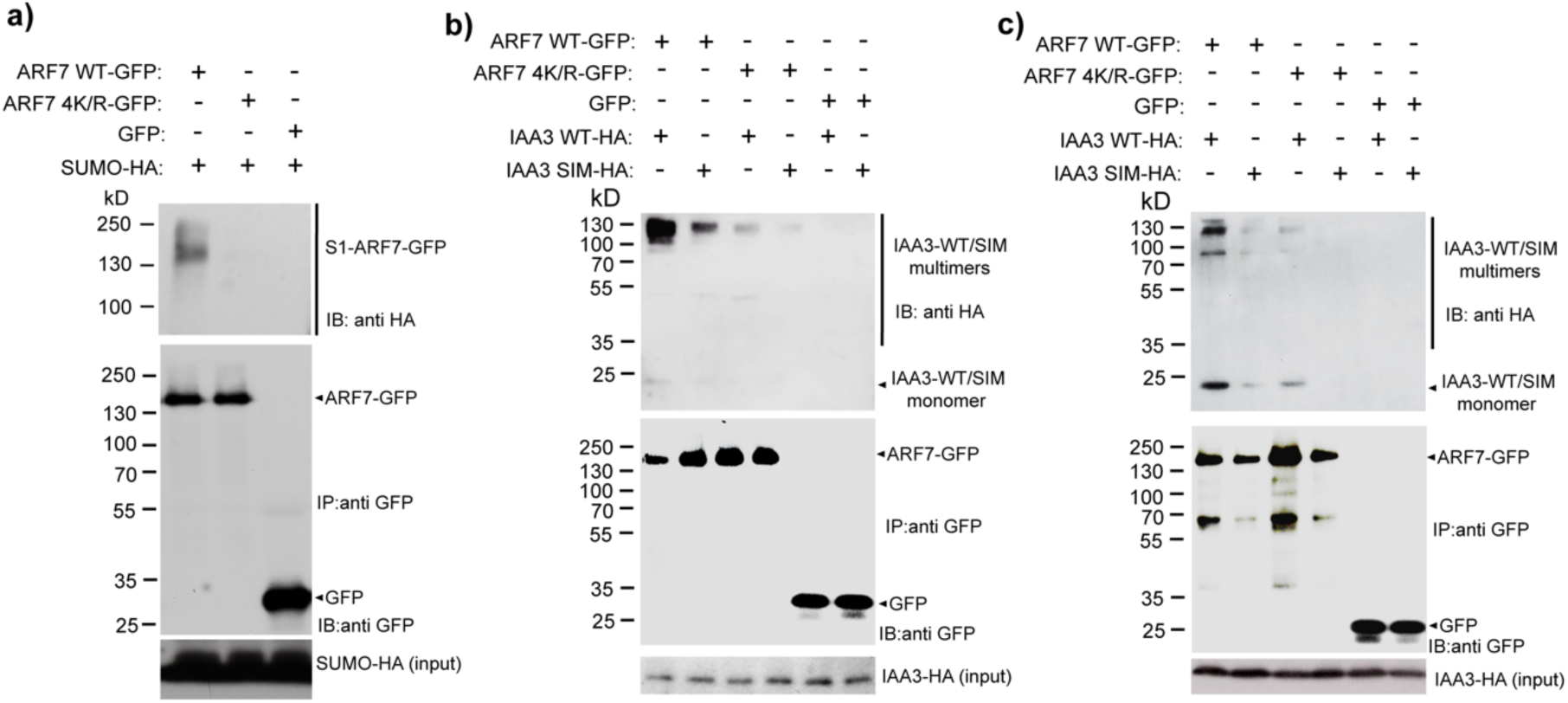
IAA3/SHY2 interaction with ARF7 is determined by the SUMO-SIM interaction module. (a) ARF7 is SUMOylated in planta as indicated in transient assays in N. benthamiana leaves co-expressing ARF7 WT-GFP or ARF7 4K/R-GFP and SUMO-HA. Multimerization status of IAA3/SHY2 is not a determinant of its interaction with ARF7 as both (b) exclusively multimeric form of IAA3 and (c) a heterogeneous population of monomeric and multimeric forms of IAA3 remains able to interact with ARF7 WT through the SUMO-SIM interaction module

**Supplementary figure S10.**
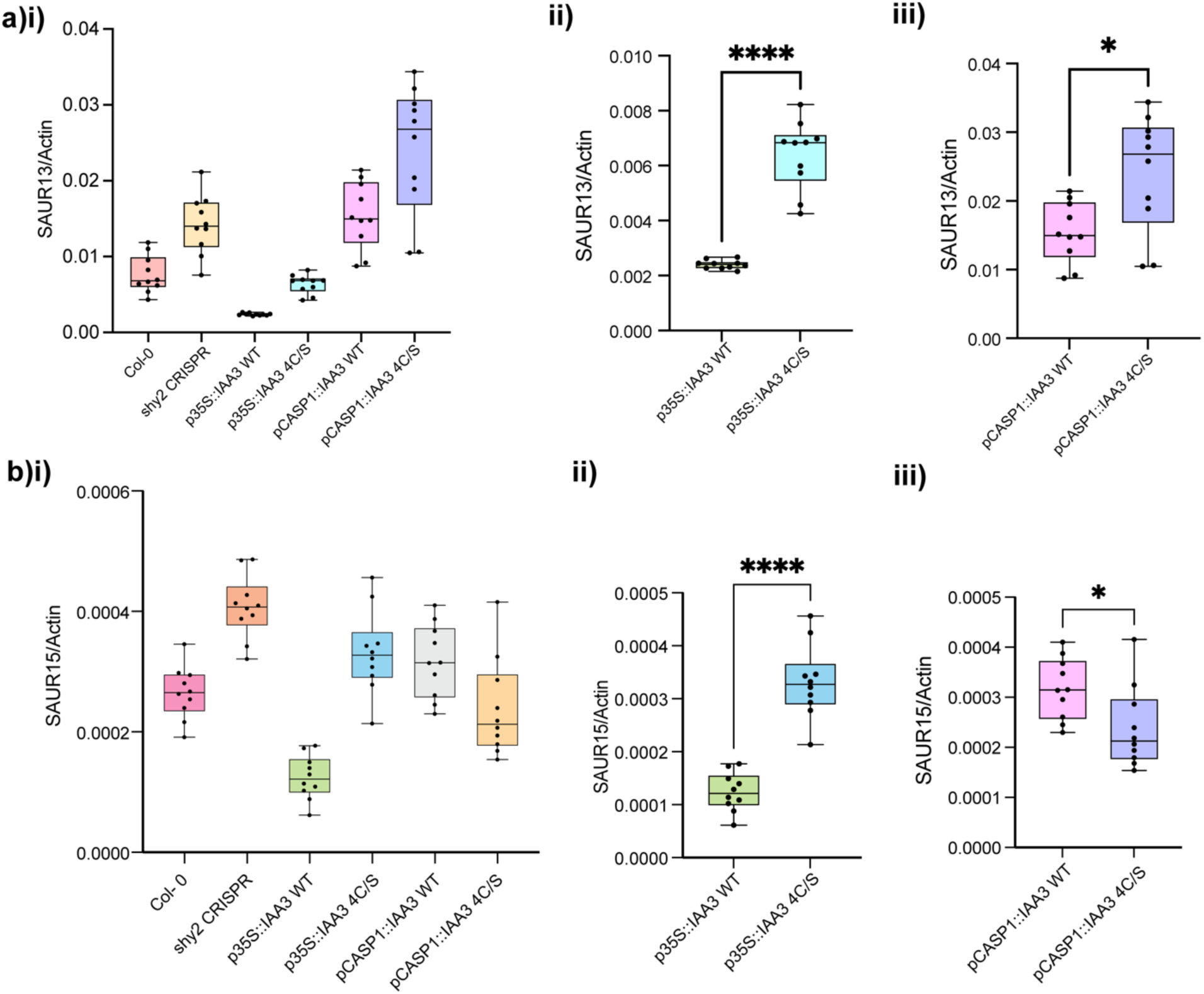
Expression profile of representative *SAUR* genes that are downstream targets of SHY2/IAA3 in the absence of exogeneous auxin treatment. **(a)** SAUR13 **(b)** SAUR15. qRT-PCR analysis was done using the absolute quantification method and the points within the box plots indicates replicates of individual experiments done using at least two independent lines for IAA3-WT and IAA3-4C/S expressing transgenics. ns, not significant, *P ≤ 0.05;***P≤ 0.001 ****P ≤ 0.0001

**Supplementary figure S11.**
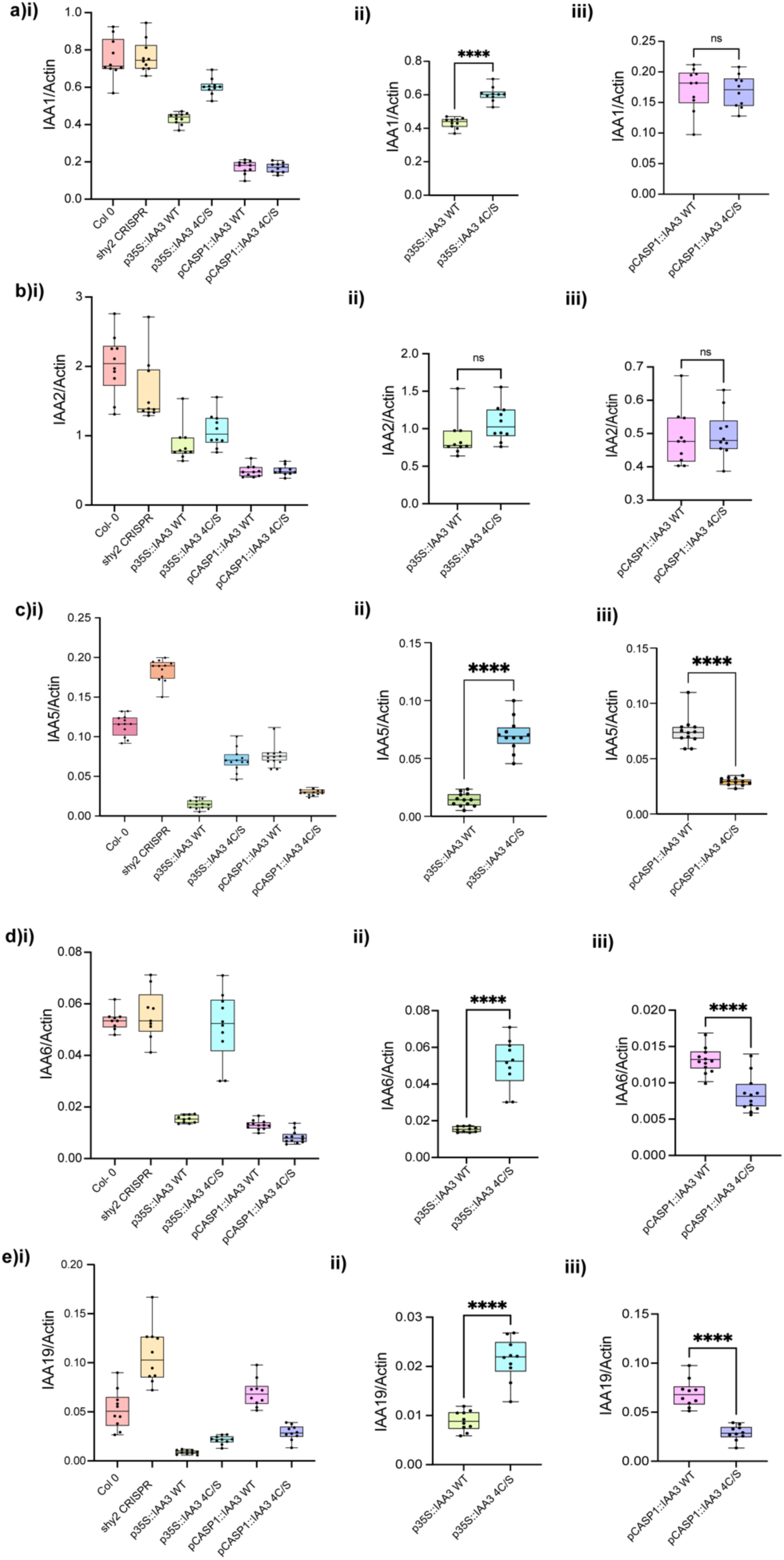
Expression profile of representative Aux/IAAs that are downstream targets of SHY2/ IAA3 in the absence of exogeneous auxin treatment. **(a)** IAA1 **(b)** IAA2 **(c)** IAA5 **(d)** IAA6 **(e)** IAA19. qRT-PCR analysis was done using the absolute quantification method and the points within the box plots indicates replicates of individual experiments done using at least two independent lines for IAA3-WT and IAA3-4C/S expressing transgenics. ns, not significant, *P ≤ 0.05;***P≤ 0.001 ****P ≤ 0.0001.

**Supplementary figure S12.**
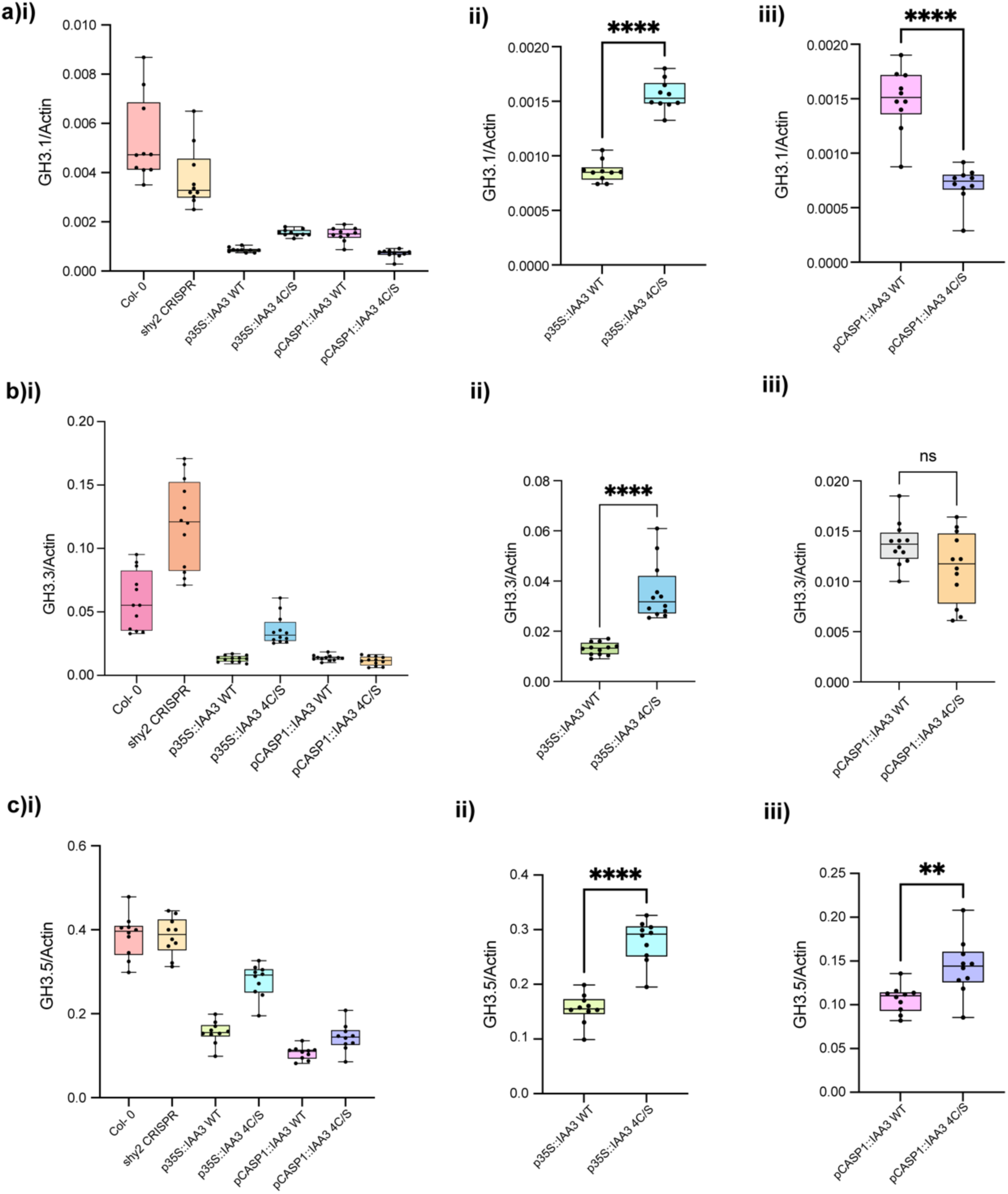
Expression profile of representative *Gretchen Hagen* (GH) genes that are downstream targets of SHY2/ IAA3 in the absence of exogeneous auxin treatment. **(a)** GH3.1 **(b)** GH3.3 **(c)**GH3.5. qRT-PCR analysis was done using the absolute quantification method and the points within the box plots indicates replicates of individual experiments done using at least two independent lines for IAA3-WT and IAA3-4C/S expressing transgenics. ns, not significant, *P ≤ 0.05;***P≤ 0.001 ****P ≤ 0.0001.

**Supplementary figure S13.**
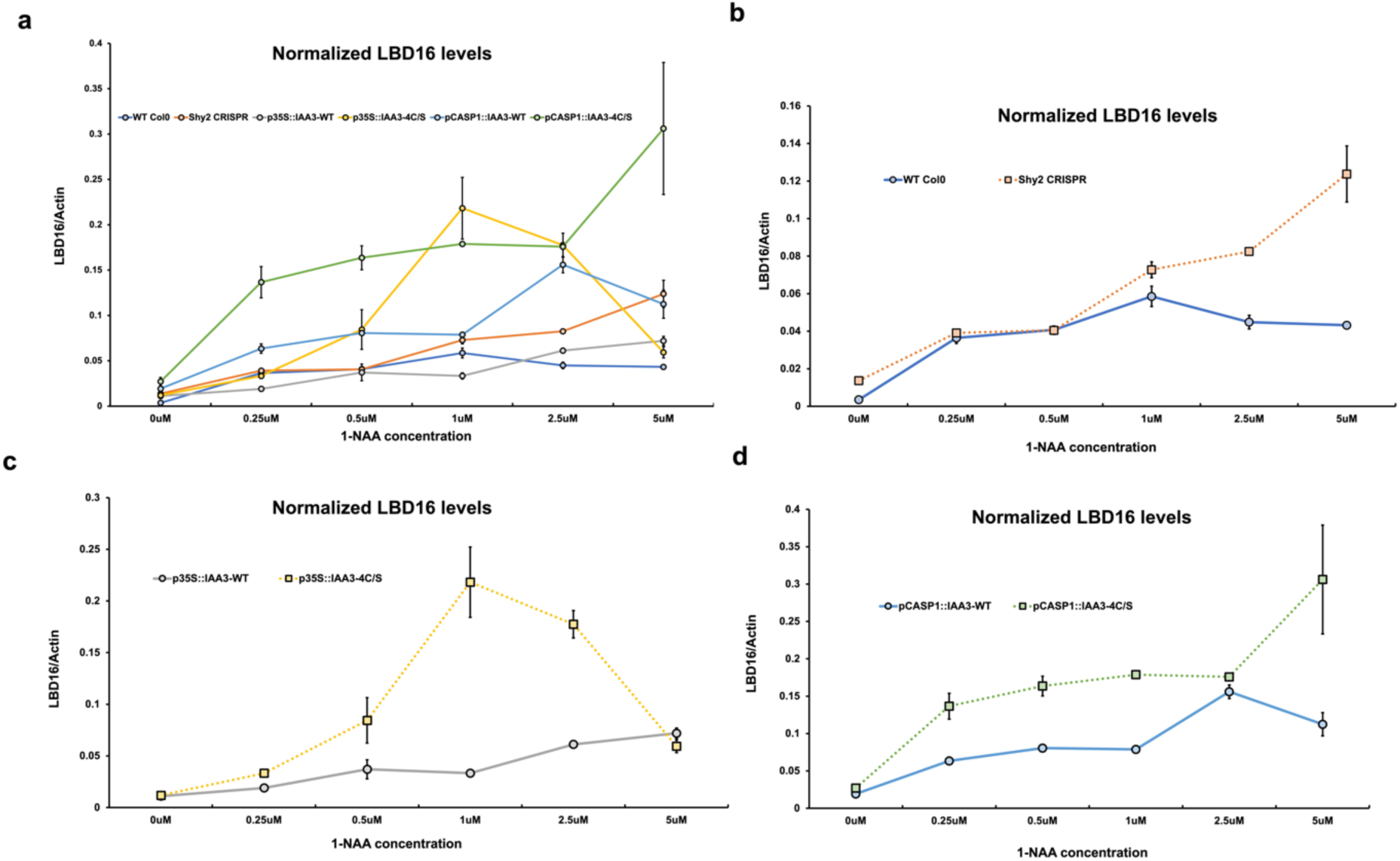
Inducibility of LBD16 on exogeneous auxin (1-NAA) treatment in IAA3WT and IAA3-4C/S expressing lines. **(a)** Expression of LBD16 in WT col0, shy2 CRISPR, p35S::IAA3-WT, p35S::IAA3-4C/S, pCASP1::IAA3WT and pCASP1::IAA3-4C/S lines. Comparative analysis of the inducibility of LBD16 between (b) WT col0 and shy2 CRISPR (c) p35S::IAA3-WT and p35S::IAA3-4C/S (d) pCASP1::IAA3WT and pCASP1::IAA3-4C/S lines upon exogeneous auxin (1-NAA) treatment. Error bars indicate standard error mean for three independent replicates.

**Table. S1.** DNA constructs used for this study. List of constructs generated by either conventional restriction enzyme-based cloning or Gateway recombination cloning technologies for performing various experiments related to this study.

| Genes | Gene Constructs | Background Vectors | Experiments |
| --- | --- | --- | --- |
| WT IAA3 | MBP-WT IAA3 | pMal c5X | In vitro redox assays |
| 4C/S IAA3 | MBP-4C/S IAA3 | pMal c5X | In vitro redox assays |
| WT IAA28 | MBP-WT IAA3 | pMal c5X | In vitro redox assays |
| WT IAA13 | MBP-WT IAA3 | pMal c5X | In vitro redox assays |
| WT IAA14 | MBP-WT IAA3 | pMal c5X | In vitro redox assays |
| WT IAA19 | MBP-WT IAA3 | pMal c5X | In vitro redox assays |
| MBP | MBP | pMal c5X | Control for experiments |
| ARF7 | ARF7-GFP | pEG 103 | Co-IP and SUMO IP |
| ARF7 4K/R | ARF7<sup>4K/R</sup>-GFP | pEG 103 | Co-IP and SUMO IP |
| WT IAA3 | HA-WT IAA3 | pEG 201 | Co-IP and Redox IP |
| IAA3-SIM | HA-SIM IAA3 | pEG 201 | Co-IP |
| 4C/S-IAA3 | HA-4C/S IAA3 | pEG 201 | Co-IP and Redox IP |
| TIR1 | GFP-TIR1 | pEG 104 | Co-IP |
| TPL | TPL-myc | pGWB18 | Co-IP |
| NINJA | NINJA-myc | pGWB18 | Co-IP |
| SUMO1 | HA-SUMO1 | pEG 201 | SUMO IP |
| GFP | GFP | pEG103/pEG104 | Control for experiments |
| WT IAA3 | p35S::HA-WT IAA3 | pEG 201 | Generation of transgenic plants |
| WT IAA3 | pCASP1::HA-WT IAA3 | pEG 201 | Generation of transgenic plants |
| WT IAA3 | pGATA23::HA-WT IAA3 | pEG 201 | Generation of transgenic plants |
| 4C/S IAA3 | p35S::HA-4C/S IAA3 | pEG 201 | Generation of transgenic plants |
| 4C/S IAA3 | pCASP1::HA-4C/S IAA3 | pEG 201 | Generation of transgenic plants |
| 4C/S IAA3 | pGATA23::4C/S-WT IAA3 | pEG 201 | Generation of transgenic plants |

**Table. S2.**
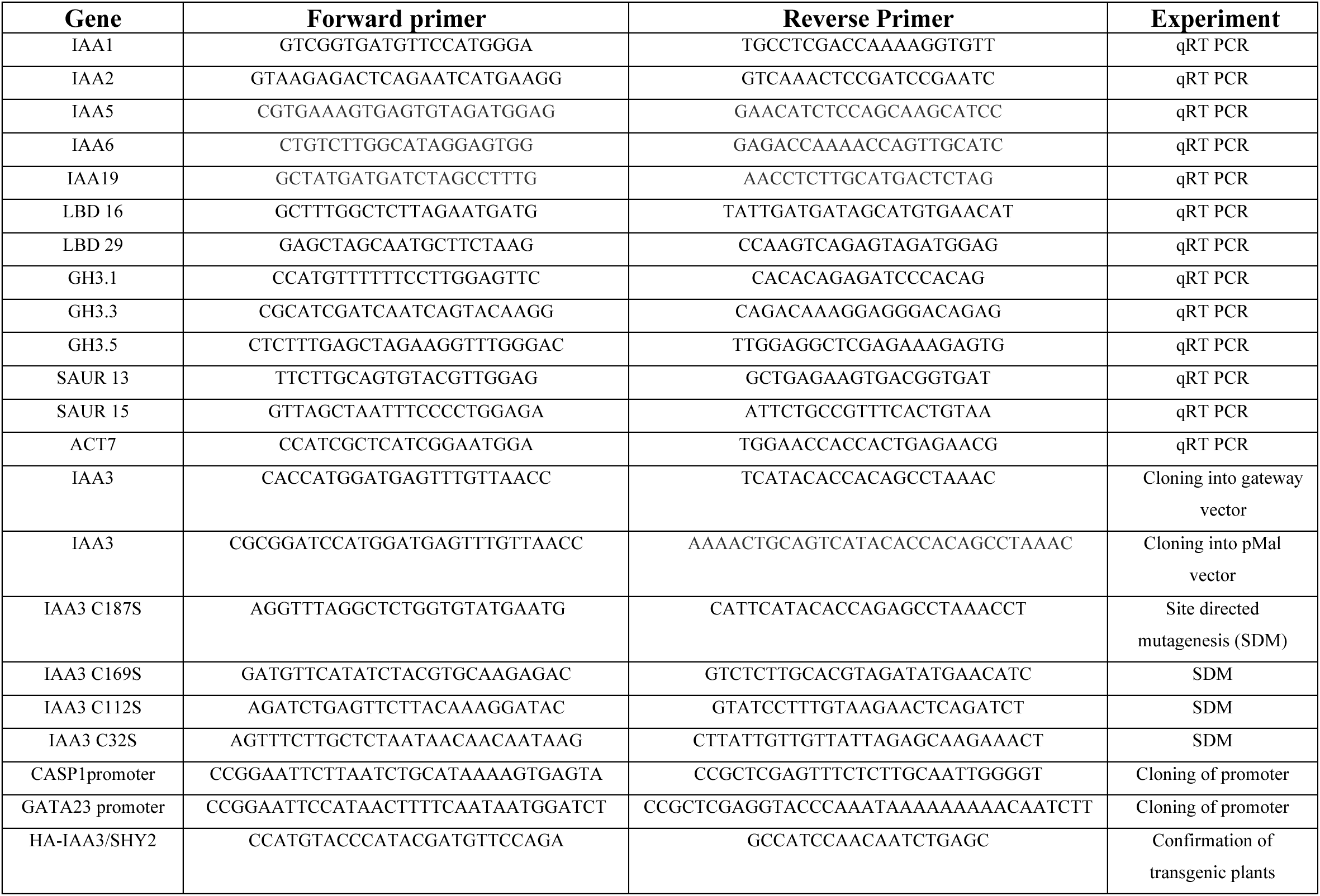
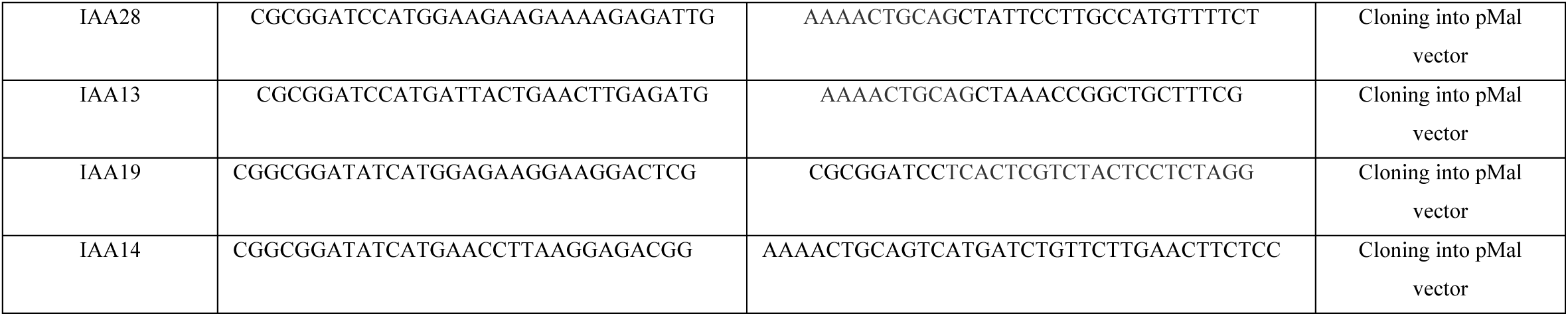
Primers used for this study. List of primers used for this study.

## Notes

### Competing Interest Statement

The authors have declared no competing interest.

